# Relative salience signaling within a thalamo-orbitofrontal circuit governs learning rate

**DOI:** 10.1101/2020.04.28.066878

**Authors:** Vijay Mohan K Namboodiri, Taylor Hobbs, Ivan Trujillo Pisanty, Rhiana C Simon, Madelyn M Gray, Garret D Stuber

**Affiliations:** The Center for the Neurobiology of Addiction, Pain, and Emotion, Department of Anesthesiology and Pain Medicine, Department of Pharmacology, University of Washington, Seattle, WA 98195, USA; Alcohol and Addiction Research Group, Department of Neurology, Neuroscience Graduate Program, University of California at San Francisco, San Francisco, CA, 94158, USA; Graduate Program in Neuroscience, University of Washington, 98195

## Abstract

Learning to predict rewards is essential for the sustained fitness of animals. Contemporary views suggest that such learning is driven by a reward prediction error (RPE) — the difference between received and predicted rewards. The magnitude of learning induced by an RPE is proportional to the product of the RPE and a learning rate. Here we demonstrate using two-photon calcium imaging and optogenetics in mice that certain functionally distinct subpopulations of ventral/medial orbitofrontal cortex (vmOFC) neurons signal learning rate control. Consistent with learning rate control, trial-by-trial fluctuations in vmOFC activity positively correlates with behavioral updating when RPE is positive, and negatively correlates with behavioral updating when RPE is negative. Learning rate is affected by many variables including the salience of a reward. We found that the average reward response of these neurons signals the relative salience of a reward, as it decreases after reward prediction learning or the introduction of another highly salient aversive stimulus. The relative salience signaling in vmOFC is sculpted by medial thalamic inputs. These results support emerging theoretical views that the prefrontal cortex encodes and controls learning parameters.

## INTRODUCTION

A simple, yet powerful model for learning that a cue predicts an upcoming reward is to update one’s predictions of future reward via a reward prediction error signal (RPE)—the difference between a received and predicted reward (Rescorla and Wagner, 1972; Schultz et al., 1997). This RPE model largely explains the response dynamics of midbrain dopaminergic neurons (Eshel et al., 2015; Mohebi et al., 2019; Schultz et al., 1997)—the main identified neural system responsible for reward prediction learning (Chang et al., 2016; Lee et al., 2020; Steinberg et al., 2013). Even though the bulk of experimental work into the neuronal mechanisms of reinforcement learning focuses on the mesolimbic dopamine circuitry, a rich computational literature argues that the dopamine RPE circuitry cannot operate in isolation and that the brain likely contains complementary systems for learning (Schweighofer and Doya, 2003; Soltani and Izquierdo, 2019; Wang et al., 2018). For instance, reinforcement learning algorithms contain parameters such as learning rate that also need to be learned. Since the net magnitude of learning due to an RPE is the product of the RPE and a learning rate for the reward, optimally tuning the learning rate (amount by which RPE updates reward prediction) can be highly beneficial to adapt learning to one’s environment (Behrens et al., 2007; Iigaya, 2016; Schweighofer and Doya, 2003; Soltani and Izquierdo, 2019; Wang et al., 2018). One of the earliest models to suggest that the learning rate is itself dynamic was the Pearce-Hall associative learning model, which proposed that learning rate is proportional to the absolute magnitude of the previous RPE (Pearce and Hall, 1980). In this model, the absolute magnitude of the previous trial’s RPE is generally referred to as the salience of the reward. Thus, an unpredicted reward has high salience and learning rate, but a predicted reward has low salience and learning rate. Learning rate has also been proposed to depend on a relative salience signal, which compares the salience of a given environmental stimulus against other stimuli (Bower and Trabasso, 1964; Downing, 1968). Other models propose that learning rates depend on the variability of previously experienced RPEs or estimates of uncertainty in the environment (Behrens et al., 2007; Courville et al., 2006; Iigaya, 2016; Preuschoff and Bossaerts, 2007; Soltani and Izquierdo, 2019). Such learning of learning rate is a special case of the learning of parameters for reinforcement learning—previously referred to as “meta learning” or “meta-reinforcement learning” (Schweighofer and Doya, 2003). Adaptive learning rates are also a consequence of more recently proposed second-order learning mechanisms operating on top of an initial model-free reinforcement learning mechanism (also referred to as “meta-reinforcement learning”)(Wang et al., 2018). A common consequence of all the above learning algorithms is that the learning adapts to the properties and statistics of its environment.

Since prior research has demonstrated causal roles for the orbitofrontal cortex (OFC) in reward learning and adaptation (Constantinople et al., 2019; Jones et al., 2012; Miller et al., 2018; Namboodiri et al., 2019; Wilson et al., 2014), we hypothesized that OFC neural activity in response to a reward controls learning rate. This central hypothesis was based on our previous observation that the suppression of reward responses in vmOFC neurons was sufficient to reduce the rate of behavioral reward prediction learning in the presence of RPEs (Namboodiri et al., 2019). Here, we first tested whether trial-by-trial variability in vmOFC activity represents learning rate. Next, we used longitudinal activity tracking of the same vmOFC neuronal population across multiple contexts to test whether their reward responses adapt across these contexts consistent with relative salience signaling. Lastly, we tested whether reward response adaptation in vmOFC is mediated, at least in part, by one of its major inputs from medial thalamus.

## RESULTS

### Theoretical prediction for learning rate control

We first developed the following theoretical approach to identify a learning rate control signal. The update in reward prediction after experiencing an RPE is the product of the RPE and the learning rate (**Fig 1A**). In other words, the learning rate amplifies the effect of an RPE on the subsequent reward prediction update. Thus, if a particular set of neurons control learning rate, variability in their activity (i.e. in the control signal) should produce corresponding variability in reward prediction update. Importantly, while RPE can be positive or negative on a trial, the learning rate is a positive number. Hence, on a positive RPE trial in which reward prediction update is positive, higher learning rate should produce an even more positive reward prediction update on the next trial. This is because a higher learning rate amplifies the effect of the positive RPE. Therefore, the correlation between a learning rate control signal and the subsequent reward prediction update should be positive on positive RPE trials (**Fig 1A, B**). However, on a negative RPE trial in which reward prediction update is negative, higher learning rate should produce an even more *negative* reward prediction update on the next trial. This is because a higher learning rate amplifies the effect of the negative RPE. Thus, the correlation between a learning rate control signal and the subsequent reward prediction update should be *negative* on negative RPE trials (**Fig 1A, C**).

**Fig 1:**
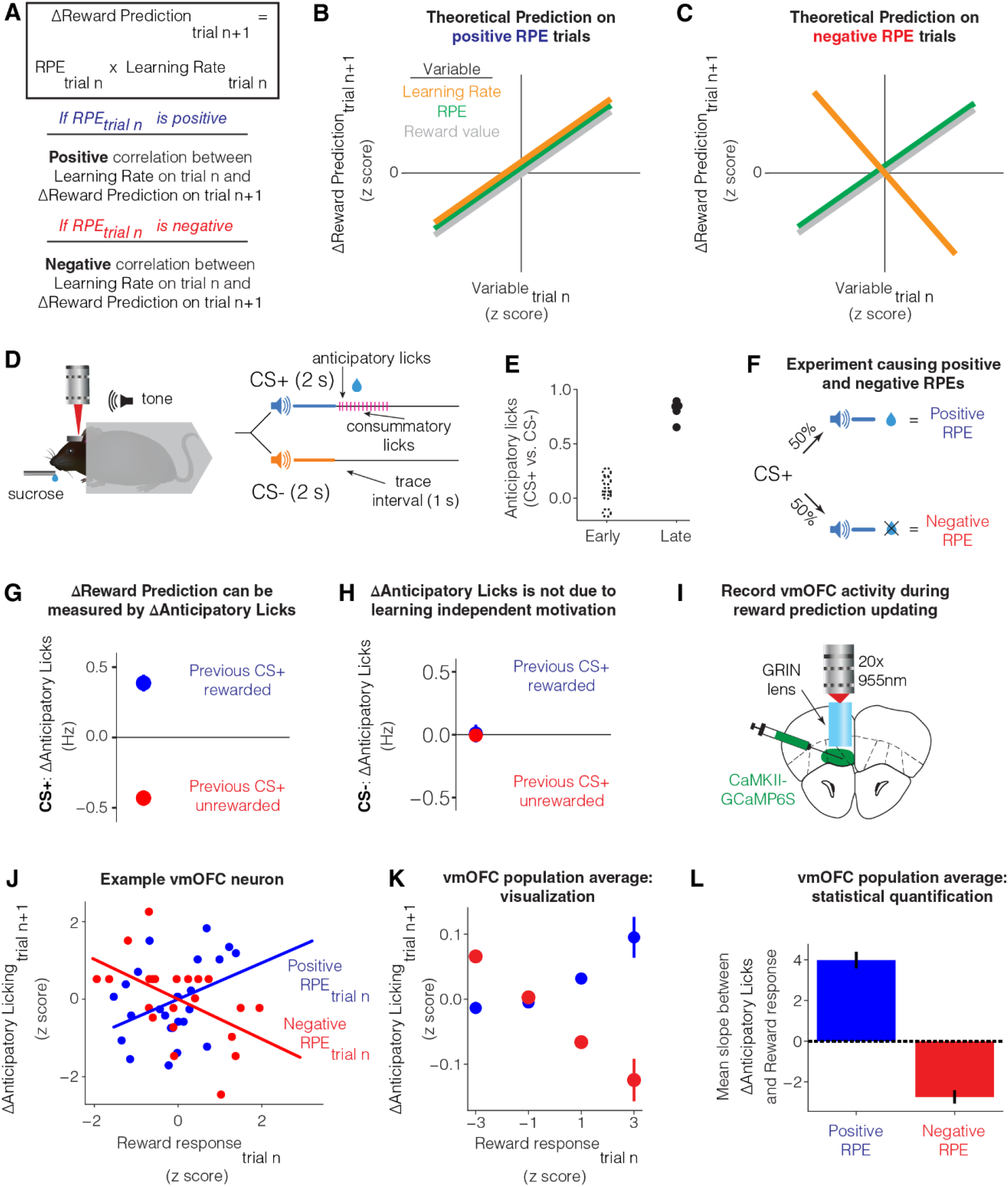
Trial-by-trial fluctuations in vmOFC reward responses reflect learning rate control. **A.** Schematic showing that the update in reward prediction on a trial from the previous trial is the RPE on the previous trial multiplied by the learning rate on the previous trial. Since learning rate has a positive correlation with RPE (Pearce and Hall, 1980), the above relationship implies that the sign of the dependence between reward prediction update and learning rate on a trial is the same as the sign of the RPE on that trial. **B.** Theoretical prediction for the dependence between trial-by-trial update in reward prediction on consecutive trials, and trial-by-trial fluctuations in different variables, for positive RPE trials. When RPE is positive, an increase in learning rate would cause an increase in reward prediction. When RPE is positive, an increase in either reward value or RPE would cause an increase in reward prediction. **C.** Theoretical prediction for the dependence between trial-by-trial update in reward prediction on consecutive trials, and trial-by-trial fluctuations in different variables, for negative RPE trials. When RPE is negative, an increase in learning rate would cause a *decrease* in reward prediction. This is because the learning rate modulates the magnitude of the reward prediction update in the direction of the RPE. On the other hand, when RPE is negative, an increase in either reward value or RPE would still cause an increase in reward prediction. This is because the slope for the relationship between RPE or reward value and reward prediction update, is the learning rate, a positive quantity. **D.** Differential trace conditioning task in headfixed mice (Namboodiri et al., 2019; Otis et al., 2017). **E.** Behavior early and late in learning. Early session was defined as the first day of learning and late session as the day when anticipatory licking in response to CS+ was high and stable (Materials and Methods) (Namboodiri et al., 2019). Cue discrimination was measured as two times the area under a receiver operator characteristic curve between lick counts after CS+ and lick counts after CS−, minus one (Materials and Methods) (Namboodiri et al., 2019). **F.** Schematic showing that in a session with reward probability of 50%, RPE will be positive on rewarded trials and negative on unrewarded trials. **G.** The change in anticipatory licking on consecutive CS+ trials (potentially with interceding CS− trials) is reliably positive after rewarded CS+ trials (positive RPE) and negative after unrewarded CS+ trials (negative RPE) (n=34 sessions from n=12 imaging mice). Thus, update in anticipatory licks can be used to estimate update in reward prediction. **<u>See Table S1 for all statistical results in the manuscript for all figures, including all statistical details and sample sizes.</u>** **H.** A potential concern is that receiving reward on a CS+ trial might increase general motivation to lick on the next trial, independent of reward prediction learning. If true, CS− trials immediately following a rewarded CS+ trial should show higher licking compared to CS− trials immediately following an unrewarded CS+ trial. This panel plots the update in anticipatory licking on CS− trials based on whether the immediately preceding CS+ trial was rewarded or unrewarded. The lack of licking update shows that the effect in **G** is not a learning independent motivation signal. **I.** Schematic showing the recording of vmOFC activity using two-photon microendoscopic calcium imaging. **J.** Data from an example neuron showing the dependence between trial-by-trial update in anticipatory licking on CS+ trials, and trial-by-trial fluctuations in response on rewarded trials (positive RPE) and unrewarded trials (negative RPE). The lines show the best fit regression in each condition. The observed relationship is as expected if vmOFC controls learning rate on a trial-by-trial basis. **K.** Z-scored, pooled, and binned data across all vmOFC neurons to visualize the dependence between trial-by-trial response fluctuations and licking update for the population of vmOFC neurons on positive and negative RPE trials. Each neuron’s data were z-scored separately for each axis, all z-scored data were then pooled, and binned into the four bins shown in the plot. Error bars are standard error of the mean. These data are shown purely for an intuitive visualization of the average relationship between these variables in the vmOFC population. **L.** Statistical quantification of the average slope between reward response on a trial and licking update on the next trial across all neurons on both positive and negative RPE trials. No z-scoring was performed here to avoid assigning an equal weight to neurons with low or high trial-by-trial variability in responses.

In contrast with the above predictions, the correlation between RPE on a trial and the subsequent reward prediction update should always be positive regardless of the sign of RPE. This is because the slope of this correlation is the learning rate, a positive quantity. Similarly, because RPE is received reward minus predicted reward, the correlation between the value of received reward on a trial and the reward prediction update on the next trial is also the learning rate, and hence always positive. In other words, unlike learning rate, an increase in RPE or reward value on a trial should always cause an increase in reward prediction on the next trial. We used this discriminative test to identify whether vmOFC activity abides by learning rate control.

The above predictions for learning rate control are also true for a signal that controls the absolute magnitude of the RPE. Indeed, absolute magnitude of the RPE has itself been hypothesized to be a driver of learning rate (Pearce and Hall, 1980). Hence, we will consider a signaling of the absolute magnitude of the RPE as another model of learning rate control for now and will consider this possibility later in the manuscript.

### Trial-by-trial and average activity in vmOFC outcome responses reflects learning rate control

To investigate the above test for learning rate control, we behaviorally measured reward prediction learning on a discriminative Pavlovian trace conditioning task in head-fixed mice (Namboodiri et al., 2019; Otis et al., 2017) (**Fig 1D**). In this task, an auditory cue paired with a delayed reward (labeled CS+ for conditioned stimulus predictive of reward) and another auditory cue paired with no reward (labeled CS−) were randomly interleaved across trials (**Fig 1D**). Once learned, mice show anticipatory licking following CS+, but not CS− (**Fig 1E**). Thus, the anticipatory licking is a behavioral proxy for reward prediction. We next verified that, consistent with reward prediction, anticipatory licking updates positively after positive RPE and negatively after negative RPEs. To consistently induce positive and negative RPEs, we reduced the probability of reward following CS+ to 50% (**Fig 1F**). In this session, reward receipt following CS+ must induce a positive RPE (as received reward is larger than the predicted reward) and reward omission following CS+ must induce a negative RPE (as received reward is less than the predicted reward). We measured the update in anticipatory licking due to positive and negative RPEs. We found that if a given CS+ trial is rewarded (i.e. positive RPE), animals increased their anticipatory licking on the next CS+ trial (may occur after interleaved CS− trials) (**Fig 1G**). Similarly, if a given CS+ trial is unrewarded (i.e. negative RPE), animals reduced their anticipatory licking on the next CS+ trial (**Fig 1G**). Thus, consistent with reward prediction update, anticipatory licking increases after a positive RPE and decreases after a negative RPE.

However, it may be possible that such updating is instead driven by learning independent changes, such as the motivation to lick. If so, after a rewarded CS+ trial, an increased motivation to lick should result in an increase in anticipatory licking even on the subsequent CS− trial. To test this possibility, we identified CS− trials that immediately followed CS+ trials and assayed whether anticipatory licking on these CS− trials depended on the outcome of the previous CS+ trial. We found that the update in CS− anticipatory licking did not depend on whether the previous CS+ trial was rewarded (**Fig 1H**). Thus, an update in anticipatory licking across CS+ trials provides a behavioral measure of reward prediction update.

We next tested whether the outcome responses of vmOFC neurons is consistent with learning rate control. To measure activity in a large number of individual vmOFC neurons during reward prediction learning, we used two-photon microendoscopic calcium imaging (Namboodiri et al., 2019) (**Fig 1I**). We found that the reward responses of vmOFC neurons (measured within 3 s after reward delivery) showed positive correlations with the subsequent update in CS+ anticipatory licking on positive RPE trials (**Fig 1J-L; <u>see Table S1 for a compilation of all statistical results including all details and sample sizes for all figures</u>**). We further found that the reward omission responses of vmOFC neurons (measured within 3 s after reward omission) showed negative correlations with the subsequent update in CS+ anticipatory licking on negative RPE trials (**Fig 1J-L**). Thus, these results are consistent with learning rate control and rule out the alternative possibilities of vmOFC responses controlling RPE or reward value. This is because these models predict a positive correlation of neural responses with reward prediction update on negative RPE trials. These results further rule out vmOFC reward responses controlling learning independent factors such as motivation. Mathematically, learning independent factors can be treated as a baseline shift independent of RPE in the equation shown in **Fig 1A** (i.e. ΔReward Prediction = RPE*learning rate + learning independent factors). Thus, the slope between such learning independent factors and reward prediction update should be independent of the RPE. In other words, an increase in learning independent factors such as motivation should produce more anticipatory licking on the next trial even if the current trial has a negative RPE. Thus, the above results are also inconsistent with an encoding of motivation. Finally, we previously demonstrated that vmOFC outcome responses causally control behavioral updating (Namboodiri et al., 2019). Specifically, we observed that inhibition of vmOFC outcome responses impairs CS+ anticipatory licking update following RPE (replotted in Fig S1). Together, these results strongly support the conclusion that vmOFC outcome responses control learning rate.

We next tested whether the average activity of vmOFC neurons abides by additional predictions of learning rate signaling. For convergence of reward prediction learning in a stationary environment, theoretical reinforcement learning models require the average learning rate to reduce over time (Iigaya, 2016; Schweighofer and Doya, 2003; Sutton and Barto, 1998). Thus, a neural signal for learning rate should reduce over the course of initial learning when reward probability following CS+ was stably maintained at 100%. Consistent with this, we found that the overall response of vmOFC neurons to reward receipt reduced after learning (**Fig 2A-C**).

**Fig 2:**
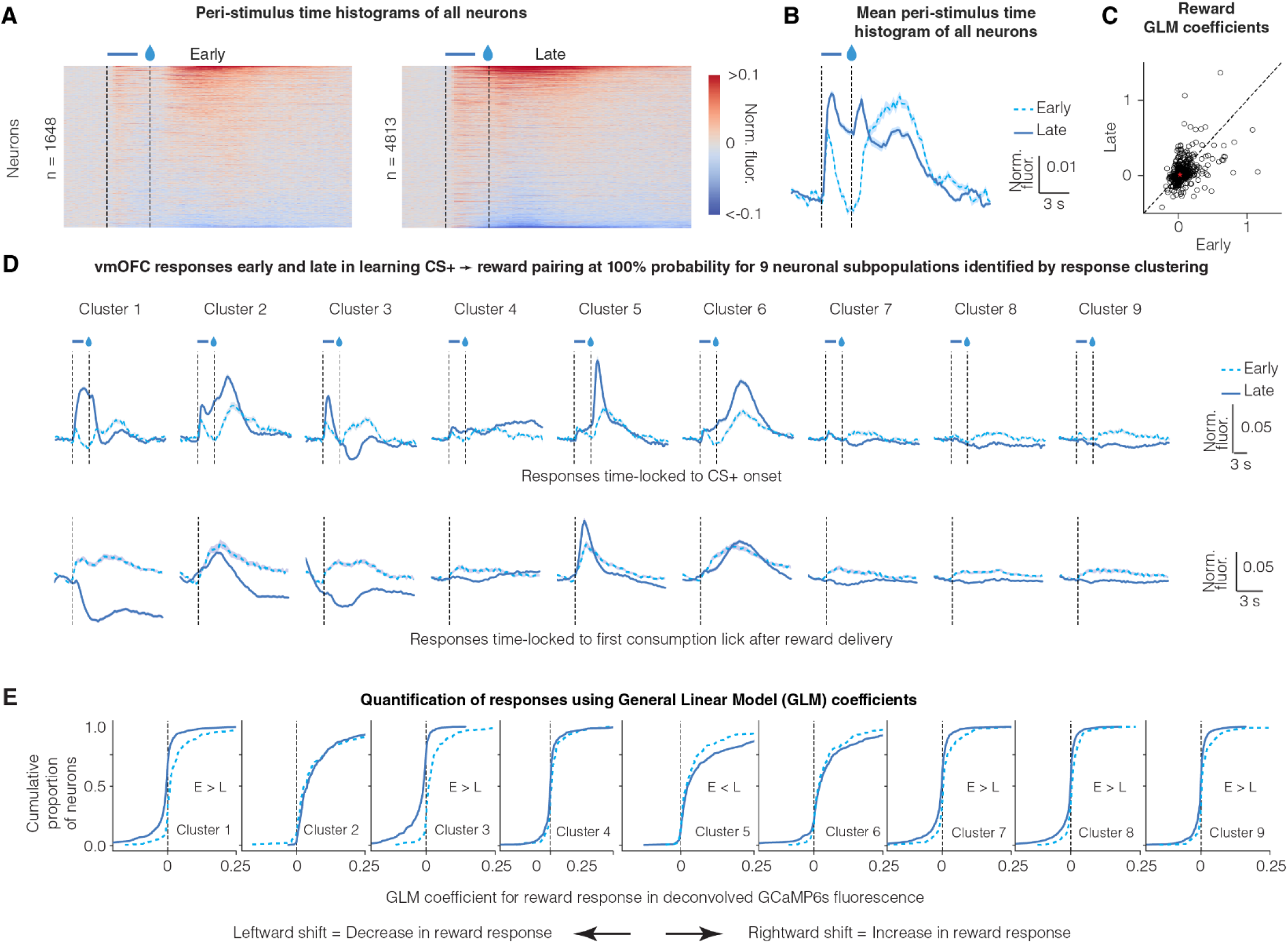
Reward responses of some vmOFC neuronal subpopulations reduce after reward prediction learning. **A.** Peri-Stimulus Time Histograms (PSTHs) of GCaMP6s fluorescence early and late in learning for all vmOFC neurons expressing CaMKIIα (excitatory neurons). All recorded neurons from both timepoints are shown. **B.** Average PSTH across all recorded neurons early and late in learning **C.** General Linear Model (GLM) coefficients for reward response early and late in learning from those neurons that were longitudinally tracked between these sessions (Materials and Methods) (Namboodiri et al., 2019) (n = 1,590 tracked neurons). We performed the GLM analyses on deconvolved fluorescence traces to remove lick response confounds and the slow decay of GCaMP6s dynamics (Namboodiri et al., 2019). The dashed line is the identity line where responses early and late are equal. The average response early and late is indicated by the red asterisk. On an average, the neuronal response to reward reduces significantly after reward prediction learning. **D.** PSTHs of GCaMP6s fluorescence early and late in learning for nine subpopulations identified using a clustering of late responses (Namboodiri et al., 2019). Each line corresponds to the average of CS+ PSTH across all neurons within a cluster (see text for rationale). The identification of neuronal subpopulations within CaMKIIα expressing vmOFC neurons using clustering algorithms, and the PSTHs late in learning, were published previously (Namboodiri et al., 2019). The top row shows responses time locked to cue onset and the bottom row shows responses time locked to reward consumption. Clusters 1 and 3 reverse the sign of their reward responses from positive (i.e. greater than baseline) early in learning to negative (i.e. less than baseline) late in learning. Error shadings correspond to confidence intervals. **E.** Cumulative distribution function for General Linear Model (GLM) coefficients for reward response early and late in learning (Materials and Methods) (Namboodiri et al., 2019). The y-axis effectively percentiles the responses shown on the x-axis. So, a reduction in reward response late in learning causes a leftward shift in the curves. These results demonstrate positive reward responses in all clusters early in learning, but negative reward responses in some clusters (1, 3, 7, 8 and 9) late in learning, showing a flip in sign.

As there is considerable heterogeneity in vmOFC neural responses, we next identified neuronal subpopulations with similar task-induced activity patterns using a clustering approach (Namboodiri et al., 2019). This approach allowed us to identify neurons that are active at similar times in the task, i.e. neuronal ensembles in vmOFC. Using this approach, we previously showed that vmOFC neurons cluster into nine different neuronal subpopulations/ensembles based on their time-locked activity profiles to the CS+ and CS− late in learning (**Fig 2D**) (Namboodiri et al., 2019). Each neuron was assigned a cluster identity based on its activity late in learning when the reward probability was 100%. Due to our ability to longitudinally track the same neurons (Namboodiri et al., 2019), we were able to evaluate the response of these neuronal clusters under many other task conditions discussed later. Among all these clusters of neurons, we found positive reward responses early in the learning of the CS+-reward association (**Fig 2D, E**). Once mice learned to predict the upcoming reward upon CS+ exposure, there was a reduction in the reward responses of many clusters (**Fig 2D, E**). Some clusters (most prominently clusters 1 and 3) even displayed a negative reward response late in learning (**Fig 2D**). These data show that the suppression of reward responses after reward prediction is specific to some neuronal subpopulations within vmOFC.

We next tested two more predictions for learning rate signaling. First, average learning rates are generally higher for negative RPE trials compared to positive RPE trials (Frank et al., 2009; Galea et al., 2015; Gershman, 2015; Kojima et al., 1996). Thus, a neural signal for learning rate should also be higher on negative RPE trials compared to positive RPE trials. Consistent with this, in the sessions in which reward probability was reduced to 50%, many clusters showed higher responses during unrewarded trials (negative RPE) compared to rewarded trials (positive RPE), including in vmOFC neurons projecting to ventral tegmental area (VTA), a midbrain regulator of learning (**Fig S2**). Second, learning rate should be higher in the presence of environmental variability (Iigaya, 2016; Schweighofer and Doya, 2003; Sutton and Barto, 1998). Consistent with this, compared to the session late in learning with 100% reward probability, the responses on rewarded trials was higher on both the 50% reward probability session or another session with unpredictable rewards in the intertrial interval, both sessions with environmental variability (**Fig S2**). Thus, the reward responses of some vmOFC neuronal subpopulations abide by the above additional predictions for learning rate signaling.

### Reward responses of specific vmOFC neuronal subpopulations signal the relative salience of a reward

As mentioned in the introduction, learning rate can be modulated by several variables. For instance, the Pearce-Hall model proposes that learning rate is modulated by the absolute magnitude of RPE (Pearce and Hall, 1980). This variable is often referred to as the salience of a reward. Thus, compared to an unpredicted reward, a predicted reward in a stationary environment should have low salience and learning rate. A signaling of salience is consistent with many of the above findings (though see Supplementary Note 2 for a detailed treatment). However, learning rate may also be modulated by the relative salience of a reward in relation to other stimuli in the environment (Bower and Trabasso, 1964; Downing, 1968). For instance, like Pearce-Hall salience, the relative salience of a reward should reduce after it becomes predicted. Unlike the absolute magnitude of the RPE however, the relative salience of an unpredicted reward should also reduce when a much more salient stimulus is introduced in the rewarding context. We next tested if the vmOFC reward responses are consistent with a signaling of the absolute magnitude of RPE or the relative salience of a reward.

To this end, we investigated whether vmOFC reward responses are suppressed in a context independent of reward prediction due to the presence of another salient stimulus. Animals typically have higher learning rates for punishments than for rewards, and for salient stimuli compared to relatively less salient stimuli (Frank et al., 2009; Galea et al., 2015; Gershman, 2015; Kojima et al., 1996; Mackintosh, 1976). For instance, prediction of highly salient aversive stimuli such as foot shocks or quinine (a bitter tastant) often occurs in single trials (Ader et al., 1972; Slotnick and Coppola, 2015). We thus hypothesized that delivering rewards in a context that also includes the delivery of salient aversive stimuli would result in a suppression of vmOFC reward responses due to a relative reduction in the salience of sucrose, *independent* of the suppression due to reward prediction (see **Supplementary Note 1** for additional reasons for the salience of quinine in these conditions). To minimize sensory confounds, we used an aversive stimulus delivered using the same sensory modality as the reward (i.e. taste). We thus intermittently and randomly (i.e. unpredictable) delivered drops of either sucrose or high concentration quinine (1.5-2.5 mM) in a 3:1 ratio to headfixed mice (**Fig 3A**) (sucrose-quinine experiment). Since the liquid deliveries were unpredictable and mostly sucrose, the animals consistently sampled the liquid to ascertain whether a given drop was rewarding or aversive. On a given trial, mice quickly suppressed their licking if the liquid was quinine, demonstrating aversion (**Fig 3B, C**).

**Fig 3:**
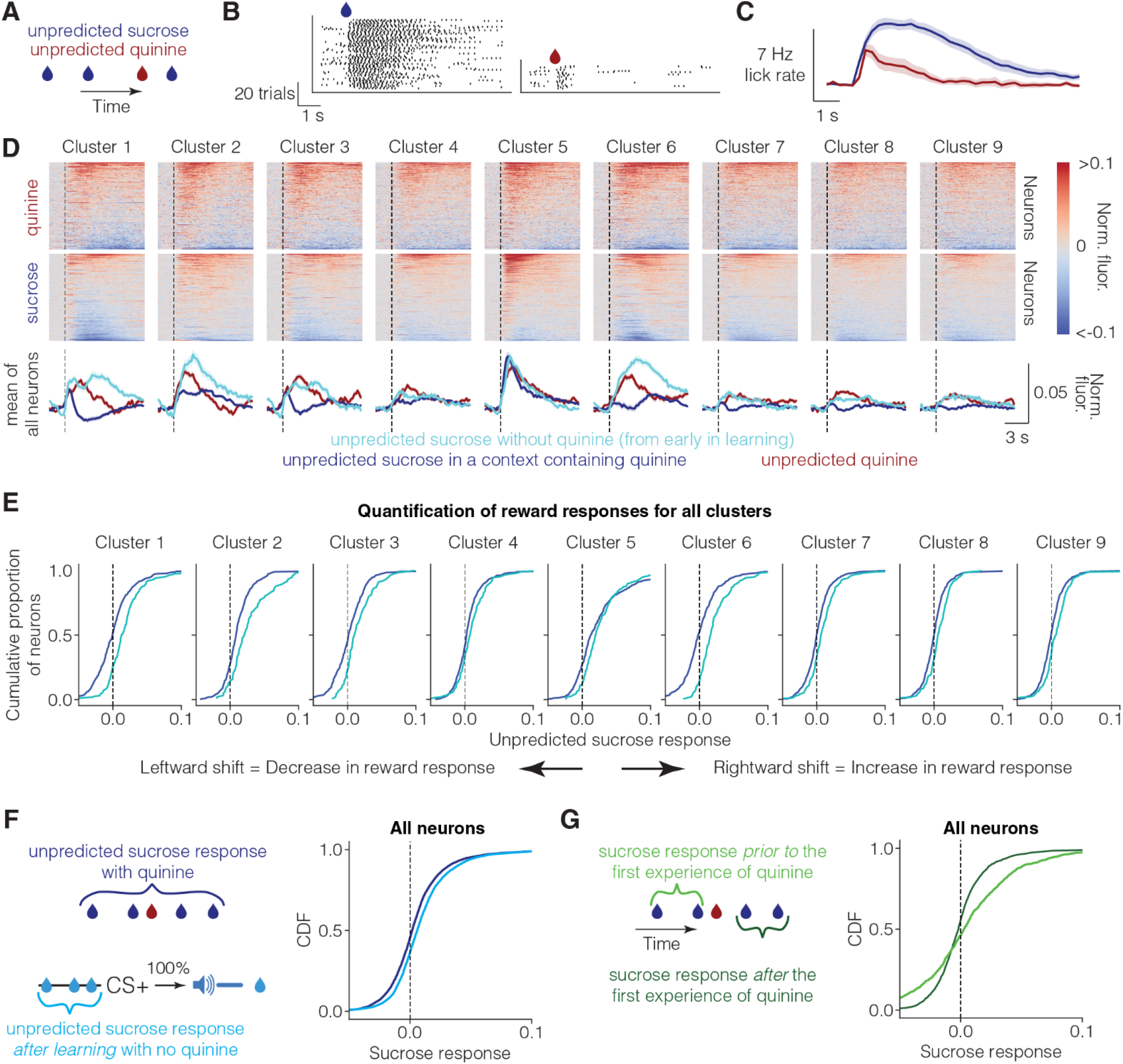
Unpredicted reward responses of vmOFC subpopulations reduce in a context containing a highly salient aversive stimulus. **A.** Schematic of unpredicted sucrose and quinine delivery. Unpredicted sucrose (10%) and quinine (1.5-2.5 mM) are delivered pseudorandomly in a 3:1 ratio (Methods). In this experiment, licks are necessary to sample the liquid. **B.** Raster plot of licking (black ticks) from an example behavioral session from one animal. Animals lick at high rates after sucrose delivery, but immediately stop licking after sampling quinine deliveries. **C.** Average lick rate across all animals and sessions (n=26 sessions from n=11 imaging mice). The histograms are time-locked to liquid delivery. **D.** The average PSTH for sucrose and quinine responses for all OFC-CaMKII neurons (n=3,716 neurons from 5 mice), aligned to the first lick after liquid delivery (i.e. initiation of consumption). The line graphs at the bottom show the average across all neurons within a cluster. The cyan line shows the average response of each cluster to reward early in learning aligned to first lick after liquid delivery (same as in Fig 2A). Responses of OFC→VTA neurons are shown in **Fig S3**. **E.** Cumulative distribution function for fluorescence responses to unpredicted sucrose in the sucrose and quinine session, and the sucrose responses early in learning. The red shadings show the clusters with the most reduction in sucrose responses due to the presence of quinine in the context. These are the same clusters as those showing suppression due to reward prediction (Fig 2**, Fig S2B**). In this case, since the response is evidently dissociated from licking (see text), we did not perform a GLM analysis. **F.** Cumulative distribution function for reward response for all vmOFC neurons for the sucrose quinine session and another control session in which unpredicted rewards were delivered during conditioning after learning (“Background” session in (Namboodiri et al., 2019)). The positive responses to unpredicted sucrose without quinine after learning further supports the positive unpredicted sucrose responses observed early in learning (shown in **D, E**). Positive unpredicted sucrose responses in the absence of quinine are also replicated in two separate cohorts in **Fig S3**. **G.** Cumulative distribution function for fluorescence responses to unpredicted sucrose in the sucrose and quinine session *prior to the first ever experience* of unpredicted quinine, and responses to unpredicted sucrose after the first experience of unpredicted quinine within the same session (i.e. same neurons). The average response of vmOFC neurons reduces after the first experience of quinine (i.e. leftward shift in curves). Note that these data only include the neurons that were recorded from the first sucrose and quinine session to only include trials before the first ever experience of quinine by the animals. Note also that the number of trials prior to the first experience of quinine is quite low (n = 2.6 trials on average), adding to the variability of responses in this condition.

Consistent with a signaling of relative salience, we found that unpredicted sucrose responses in vmOFC neurons were suppressed when delivered in a context containing quinine (**Fig 3D, E**). In comparison, reward responses early in learning (**Fig 3D, E**, **Fig 2B-D, Fig S3**), and unpredicted reward responses after learning in the absence of quinine (**Fig 3F**), were positive across all clusters. To further demonstrate the effect of quinine presentation on the response to sucrose, we compared the sucrose trials prior to the first experience of quinine against the remaining sucrose trials in the first sucrose-quinine session (**Fig 3G**). We found that the sucrose responses reduced considerably within the same session after the first experience of quinine. Overall, these results support relative salience signaling.

One potential concern with the above results could be whether the reward response adaptation due to reward prediction or quinine are correlated within the *same* neurons. To test this, we evaluated the activity of only those neurons that were longitudinally tracked across these sessions (**Fig 4**). We found that the reward responses were correlated across all three conditions in these neurons (**Fig 4**, correlations quantified in **Table S1**). Thus, the reward response adaptation due to reward prediction or the presence of quinine are correlated across the vmOFC neuronal population. We further ruled out the possibility that vmOFC reward responses simply reflect an efference copy of an arousal signal that may vary across these conditions (**Fig S4**). To this end, we tested whether changes in pupil diameter in darkness, a measure of arousal, show similar changes as those seen in vmOFC subpopulations. We found that the mean pupil diameter responses to sucrose early in learning and after 50% reward probability reduction do not correlate with the corresponding vmOFC reward responses (**Fig S4**). Nevertheless, consistent with a reduction in the relative salience of sucrose in a context containing quinine, we observed a highly dampened pupil dilation for sucrose in this context (**Fig S4**). These observations show that a simple efference copy of an arousal signal is not sufficient to explain vmOFC reward responses. Thus, consistent with a system that signals the relative salience of a reward, vmOFC reward responses reduce in two independent settings due to reward prediction or the reduction of relative salience of unpredicted rewards. Cumulatively, the reward responses observed in vmOFC rule out typically assumed reinforcement learning variables such as RPE, absolute magnitude of RPE, reward value, relative reward value and expected future value (treated in detail in **Supplementary Note 2**).

**Fig 4.**
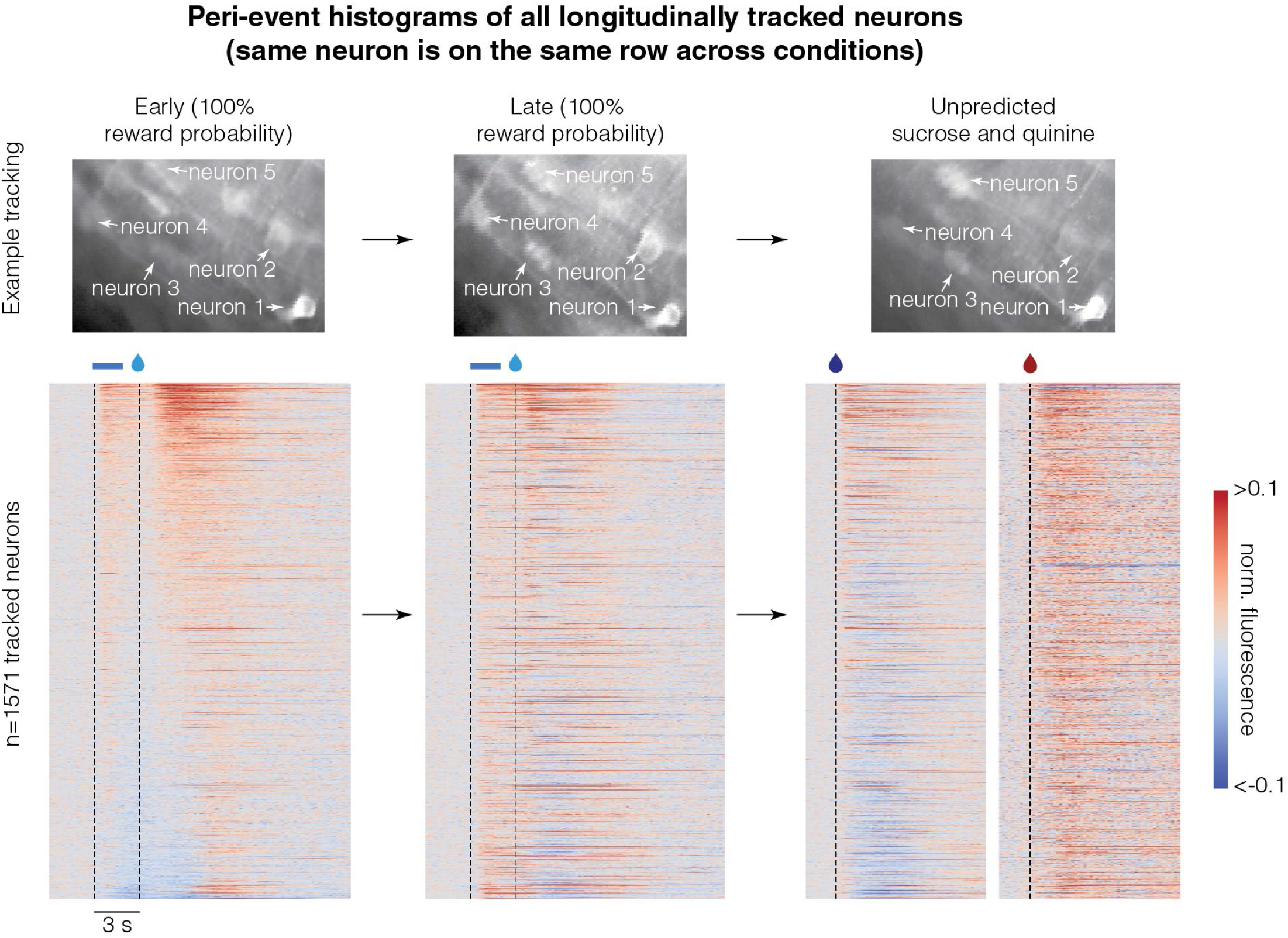
Reward responses of the longitudinally tracked neurons are correlated across three conditions: Top row shows example longitudinally tracked neurons. Here, the intensity of a pixel corresponds to activity and hence, different brightness across sessions corresponds to different activity levels. The bottom four rows show the peri-event histograms of all longitudinally tracked neurons across the four clusters showing reward response adaptation. Each row across the different conditions corresponds to the same neuron. Neurons within a cluster are sorted by their average activity early in learning. These data show that response to reward is correlated across all conditions (quantified in Table S1). For instance, the neurons that show the lowest amount of activity early in learning tend to be the neurons that show inhibitory reward responses late in learning or in the sucrose and quinine experiment (correlations quantified in Table S1). Please note that the responses of some neurons are saturated in the color map to ensure that the response patterns of most neurons are visible.

### Medial thalamic inputs to vmOFC control relative salience signaling in vmOFC

We next assessed the neuronal circuit mechanism for the relative salience signaling in vmOFC neurons. We hypothesized that reward responses in vmOFC may at least be partially controlled by inputs from medial thalamus (mThal). This is because a wide array of reward responsive regions such as basolateral amygdala, other prefrontal cortical regions, and pallidal regions project to the medial thalamus and can indirectly control vmOFC reward responses through mThal (Jankowski et al., 2013; Mitchell and Chakraborty, 2013). Further, disconnection studies have shown that interactions between mThal and OFC are necessary for reward related decision-making (Izquierdo and Murray, 2010). Despite this, whether reward responses in mThal→vmOFC input exhibit relative salience signaling and whether this input causally affects relative salience signaling in vmOFC is unknown. We first identified the anatomical locations of thalamic cell bodies projecting to vmOFC using viral (Tervo et al., 2016) and non-viral (Otis et al., 2017) retrograde tracing approaches (**Fig 5A**). The predominant thalamic structures projecting to vmOFC are the anteromedial and mediodorsal thalamic nuclei (**Fig 5B**, **Fig S5**). We then investigated the reward response plasticity of input from these regions to vmOFC. We compared unpredicted reward responses of mThal→vmOFC axons in sessions without and with quinine (**Fig 5C-E**). We found largely positive responses in these axons in response to unpredicted sucrose rewards in the absence of quinine (**Fig 5F**, **Fig S5**). These reward responses were suppressed in a session containing quinine (**Fig 5F**), showing qualitative correspondence with the reward response adaptation observed in vmOFC neuronal responses.

**Fig 5:**
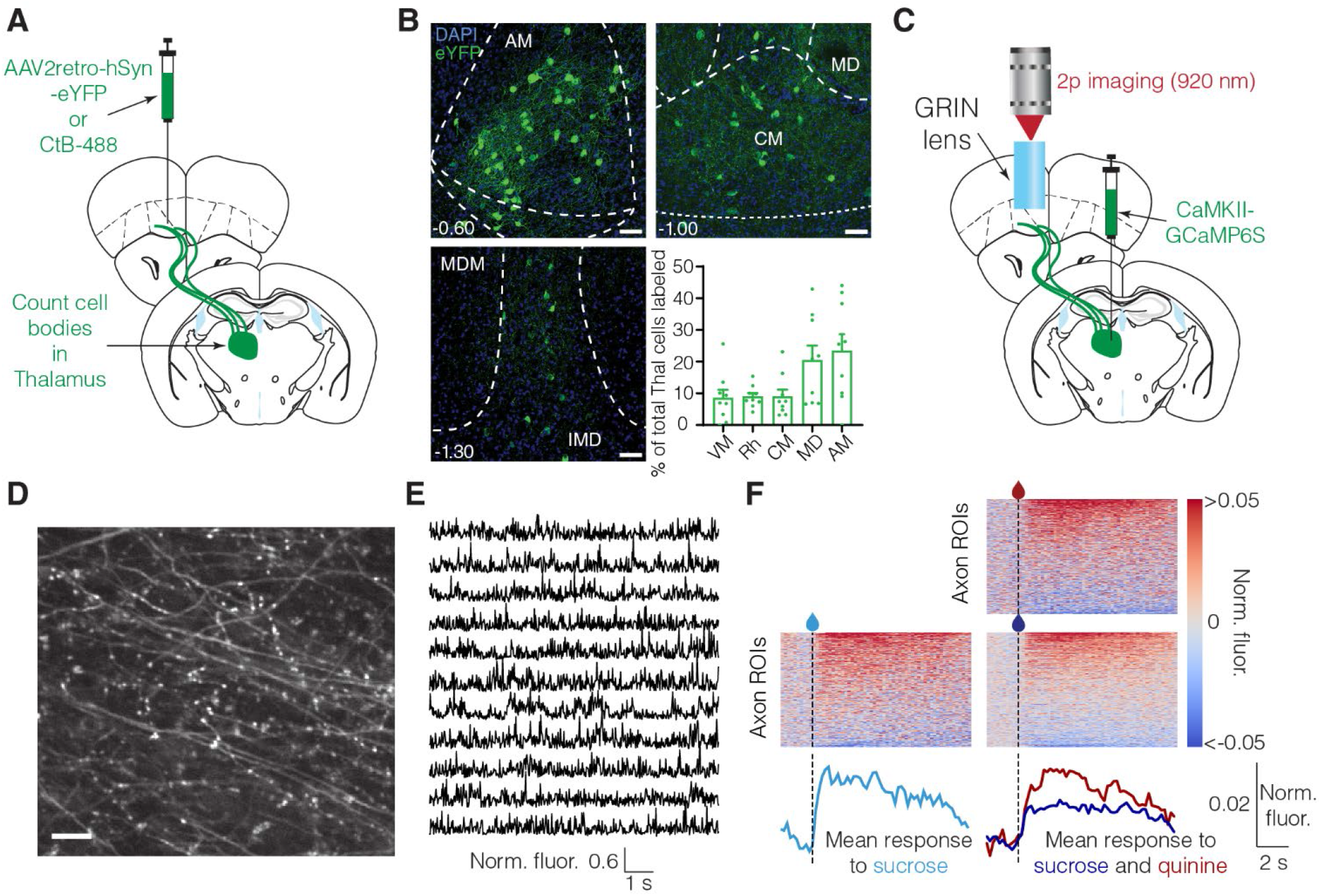
Medial thalamus (mThal) conveys reward responses to vmOFC and shows qualitatively similar reward response adaptation as vmOFC neurons. **A.** Surgery schematic for retrograde anatomical tracing, showing injections of either retrogradely traveling virus (AAV2retro) or Cholera Toxin-B (CTB). **B.** mThal cell bodies projecting to OFC counted using CTB and AAV2retro labeling. Representative images show AAV2retro expression (see **Fig S5** for CTB expression). Top 5 thalamic regions are shown. See **Fig S5** for counts in all thalamic nuclei, split by AAV2retro and CTB injections. AM: Anteromedial, MD: Mediodorsal, CM: Centromedial, Rh: Rhomboid, VM: Ventromedial. Scale bar = 50 μm. **C.** Surgery schematic for mThal axon imaging in vmOFC. **D.** Example zoomed-in mThal axon standard deviation projection image showing individual axons in vmOFC. The scale bar corresponds to 10 μm. **E.** Example mThal GCaMP traces from individual axonal regions of interest (ROIs) (Methods). **F.** Heat maps show trial-averaged responses from individual axon ROIs (*do not necessarily correspond to distinct axons,* see **Fig S5** and Methods for details and interpretation) to unpredicted sucrose alone (left) or unpredicted sucrose and quinine (right, similar experiment as Fig 3A). The bottom traces show the average responses across all segmented mThal axon ROIs aligned to the first lick after liquid delivery (dashed line). The same animals (n=3) were used under all conditions, to be directly comparable; see **Fig S5** for data from two more animals in the sucrose only condition. Mean sucrose response across the population is lower in the session with quinine compared to the session without quinine. The statistical test was applied using the mean fluorescence across all ROIs per animal as an independent measure. This adaptation is qualitatively similar to that seen in some vmOFC clusters (Fig 3).

These results suggest that mThal input might contribute to the relative salience signaling observed in vmOFC neurons. To test the causal influence of mThal input on vmOFC reward responses, we optogenetically inhibited mThal→vmOFC input after reward delivery while imaging from vmOFC neurons (**Fig 6A**). To remove light artifacts, we discarded the imaging frames during optogenetic inhibition, and evaluated reward responses right after the termination of inhibition. Since GCaMP6s responses are slow with a decay time of roughly two seconds (Chen et al., 2013), a change in activity during the inhibition will be apparent even after the inhibition for up to two seconds. We found that individual neurons showed both positive and negative modulation of activity due to mThal inhibition (**Fig 6B**). To test whether mThal input affects vmOFC reward response adaptation, we inhibited mThal→vmOFC axons in a session containing unpredicted deliveries of sucrose and quinine. We found that the reduction in sucrose responses due to the presence of quinine was significantly dampened upon mThal inhibition, in all clusters except clusters 4 and 5 (**Fig 6D**). There was also a non-selective change in quinine responses in some clusters (**Fig S6**). Two potential confounds for this experiment are that vmOFC responses may reflect the presence of light due to the LED, or that mThal→vmOFC axons may show rebound excitation after the one second inhibition. We ruled these confounds out because we observed no effect on vmOFC neurons during spontaneous inhibition of mThal→vmOFC axons in the absence of rewards, or in virus control animals without opsin expression (i.e. with LED but no inhibition) (**Fig S6**). Therefore, these results strongly support a causal role for mThal→vmOFC axons in controlling the relative salience signaling in vmOFC.

**Fig 6:**
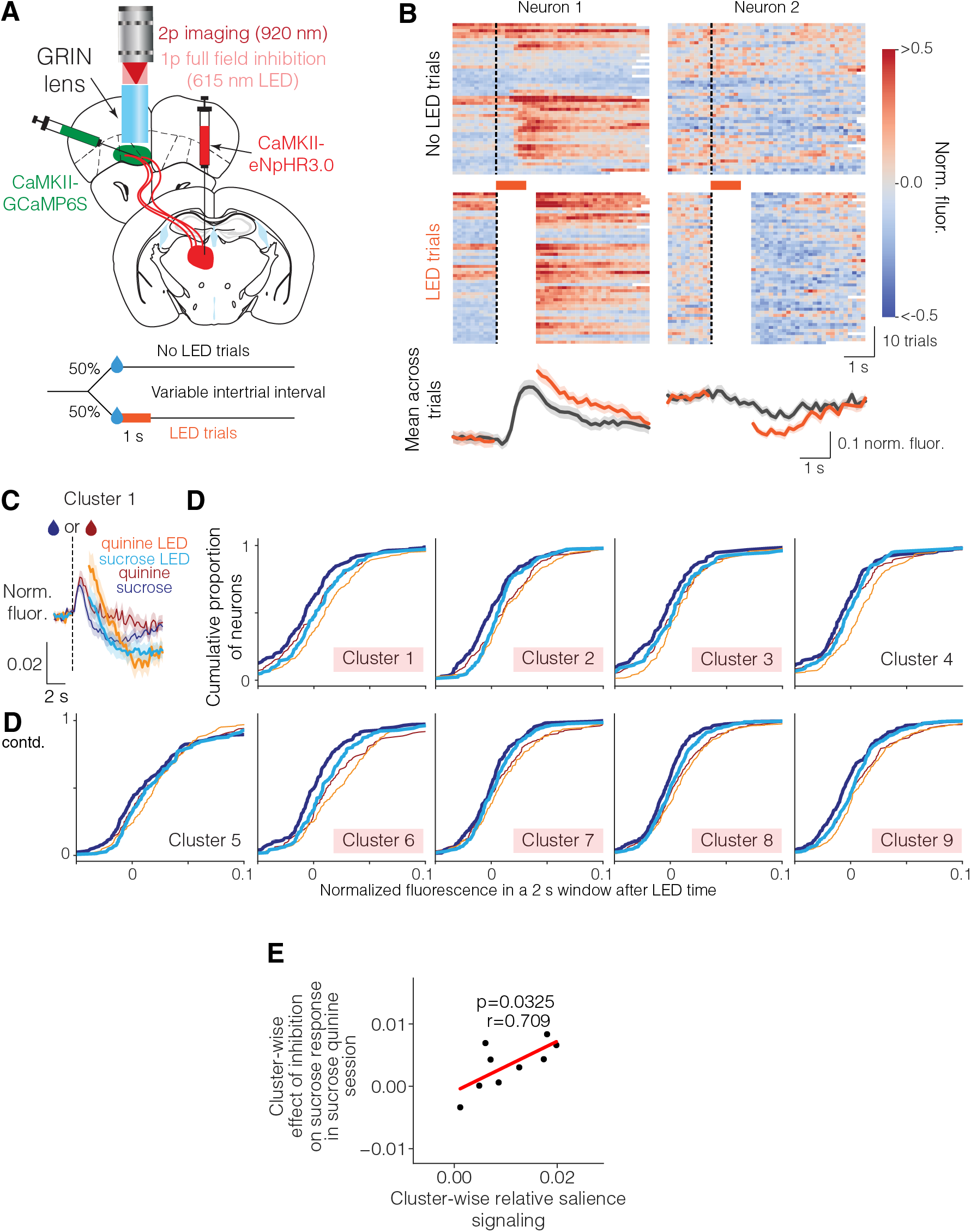
Medial thalamic input to vmOFC guides vmOFC reward response adaptation. **A.** Schematic of mThal inhibition while imaging vmOFC CaMKIIα expressing neurons. Top shows surgery schematic and bottom shows experiment schematic. **B.** Example neurons showing effect of mThal inhibition on unpredicted sucrose responses. Left neuron shows an increase in activity due to mThal inhibition, whereas the right neuron shows a decrease in activity. Frames around LED illumination were masked out (shown as white) to prevent light artifacts in imaging. **C.** PSTH of cluster 1 showing effect of mThal inhibition on sucrose and quinine responses. **D.** Empirical cumulative distribution functions showing average fluorescence with and without LED for individual neurons within each cluster for sucrose and quinine responses (n=1,645 neurons in total from 3 mice). This shows the full distribution of the population responses, with a rightward shift signifying an increase in activity. Here, the red shadings correspond to clusters showing significant mean effect on their sucrose responses. **E.** Cluster-wise relationship between relative salience signaling (measured by suppression in reward response due to the presence of quinine, Fig 3) and effect of mThal inhibition on sucrose response (i.e. change in sucrose response due to LED as shown in **D** minus the change in spontaneous response due to LED as shown in **Fig S6**). There is a strong positive correlation (∼50% explained variance).

Considering the cluster-wise variability in the effect of mThal→vmOFC inhibition on vmOFC reward responses, we next tested whether this variability is related to the variability in response adaptation across clusters. If mThal→vmOFC is responsible for controlling relative salience signaling and not just for controlling the positive reward responses (see Discussion below), the cluster-wise variability in relative salience signaling should predict the cluster-wise variability in the effect of mThal→vmOFC inhibition on vmOFC reward responses. We indeed found that the suppression of reward response due to the presence of quinine (**Fig 3**) predicts the average effect of mThal inhibition on a given vmOFC neuronal cluster/subpopulation (**Fig 6E**). Overall, these results demonstrate that the relative salience signaling in vmOFC neurons depends on mThal inputs.

## DISCUSSION

Recent studies have begun investigating OFC activity using two-photon calcium imaging (Banerjee et al., 2020; Jennings et al., 2019; Namboodiri et al., 2019; Wang et al., 2020). Using this approach to longitudinally track neuronal activity across tasks, we identified neuronal subpopulations in vmOFC that signal learning rate. We propose that the most parsimonious explanation of our major findings is a learning rate control signal (**Supplementary Note 2**). Consistent with this signal acting to control learning rate, there is significant correlation between vmOFC outcome response on a CS+ trial and the behavioral updating on the subsequent CS+ trial. Crucially, the sign of this correlation is the sign of the RPE on a trial—a strong prediction for learning rate control. Along with our previous findings that vmOFC reward responses causally controls behavioral learning based on recent reward history (Namboodiri et al., 2019), the current results provide strong evidence that vmOFC neurons act to control learning rate. Interestingly, the learning rate signaling in vmOFC is consistent with a general signaling of the relative salience of a reward across different contexts. We then show that medial thalamic inputs to vmOFC exhibit qualitatively similar response adaptation as vmOFC neurons, and causally control the relative salience signaling of specific vmOFC subpopulations. Overall, these results bolster the emerging theoretical view that the prefrontal cortex acts as a meta learning system that can adaptively control learning rate (Wang et al., 2018), in addition to representing cognitive parameters for learning such as uncertainty (Soltani and Izquierdo, 2019), confidence (Kepecs et al., 2008), surprise (Hayden et al., 2011) and volatility (Behrens et al., 2007).

In typical reinforcement learning algorithms, learning is “complete”, i.e. reward is predicted, when RPE at reward receipt becomes zero. Nevertheless, when rewards are delayed from their predictors, the resultant temporal uncertainty in the subjective estimation of the delay causes RPE to be significantly positive even after learning. Indeed, under delays less than even two seconds, midbrain dopaminergic neurons exhibit significant positive responses to a predicted reward even after extensive training (Coddington and Dudman, 2018; Cohen et al., 2012; Engelhard et al., 2019; Kobayashi and Schultz, 2008; Lee et al., 2020). Thus, it is not possible to evaluate whether a received reward was fully predicted using only the RPE signal on a trial. The reward response adaptation observed in some vmOFC subpopulations may provide an explicit signal that a delayed reward is as predicted as the reward response reverses in sign once a delayed reward is predicted. Hence, the signal from vmOFC can counteract the positive RPE signal from midbrain dopaminergic neurons and signal that a delayed reward is predicted despite the uncertainty in subjective timing. In this sense, the vmOFC reward response adaptation takes into account whether the temporal uncertainty in a delayed reward is *expected* uncertainty due to the uncertainty in estimation of the fixed delay, a computation that has been behaviorally shown to exist in rodents (Kheifets et al., 2017). A similar “brake” on learning after the prediction of a delayed reward has been assumed in computational models of the learning of reward timing (Gavornik et al., 2009; Namboodiri et al., 2015).

Numerous studies have found that the learning rate for negative RPEs is higher than the learning rate for positive RPEs (Frank et al., 2009; Galea et al., 2015; Gershman, 2015; Kojima et al., 1996). Thus, a learning rate control mechanism would produce larger responses for negative RPEs than positive RPEs. Reward responses of some vmOFC subpopulations are also consistent with this prediction since negative RPE (i.e. unrewarded) trials in a 50% reward probability session produce higher responses than positive RPE (i.e. rewarded) trials. A recent paper showed that the biased setting of learning rates on positive and negative RPE trials can provide a mechanism to learn distributions of value estimates (Dabney et al., 2020). An intriguing possibility is that the neural variability in responses to positive and negative RPEs in vmOFC neurons may reflect biases in such learning rates.

We have shown that the general role of OFC in reward learning results at least in part from the control of behavioral learning rate. Consistent with a control of behavioral learning rate by OFC, we previously found that inhibition of vmOFC→VTA neurons during the reward, but not the cue, suppresses behavioral learning based on recent reward history (Namboodiri et al., 2019). A previous study also found that inhibition of lateral OFC (likely also containing some ventral OFC) during the reward, but not cue, period of an instrumental task, affected behavioral adaptation dependent on reward history (Constantinople et al., 2019). Further, lesions of medial OFC affect learning and representation of outcomes associated with an action (Bradfield and Hart, 2020). This finding is potentially consistent with the control of learning rate, as lesion of medial OFC may cause “over-learning” of an action, thereby making it less sensitive to the expected outcome.

Careful future experiments are needed to examine the generality of these findings to different behavioral tasks and states. For instance, it has been argued that OFC function might be fundamentally different in Pavlovian versus instrumental tasks due to the differences in the mental representation of these tasks (Bradfield and Hart, 2020). Thus, it remains to be seen whether our findings would generalize to instrumental learning. Further, adaptive control of learning rate is typically studied in the context of dynamic uncertain environments. While some aspects of our task are dynamic (e.g. change in contingency while recording from the same neurons), it is necessary to test the generality of these findings in relation to the control of learning rate by expected and unexpected uncertainty (Behrens et al., 2007; Grossman et al., 2020; Iigaya, 2016; Soltani and Izquierdo, 2019). It will also be interesting to test whether OFC outcome representation in bandit tasks relate to learning rate (Costa and Averbeck, 2020). One interesting aspect of our task is that some clusters encode the outcome of a trial well past the reward was delivered (e.g. cluster 4, 6 in **Fig 1D**). Such long-lasting outcome encoding has previously been observed across different task conditions and species in the OFC (Costa and Averbeck, 2020; Hirokawa et al., 2019; Simmons and Richmond, 2008). A limitation of our study is that it was conducted under moderate water deprivation and with a specific type of reward (sucrose) and aversive stimulus (quinine). It is likely that varying the levels of water deprivation will change the behavioral salience of these liquids. Thus, careful studies are required to ascertain the generality of these findings across behavioral states and different types of rewards/punishments. Relatedly, an aberrant learning rate for drugs of abuse in OFC (e.g. higher learning rate for positive RPEs compared to negative RPEs) might partially explain the role of OFC in drug addiction (Everitt et al., 2007; Pascoli et al., 2018). If our findings generalize to these different settings, that would suggest that OFC acts more generally as a system that prioritizes currently available rewards or punishments for learning, based on the current behavioral state.

Future studies are also required to test whether relative salience signaling occurs in other brain regions. The anterior cingulate cortex has often been suggested to reflect variables required for adaptive learning rate control (Behrens et al., 2007; Hayden et al., 2011; Soltani and Izquierdo, 2019). Correlates of Pearce-Hall like salience have also been found in amygdala (Roesch et al., 2012). Nevertheless, it is unclear if these regions exhibit a strong reduction in responses to delayed, but predicted rewards and rewards presented in a context containing more salient unconditioned stimuli. Interestingly, the ventromedial prefrontal cortex, an area adjacent to orbitofrontal cortex, primarily shows lower responses to aversive outcomes than rewarding outcomes at the single neuron level (Monosov and Hikosaka, 2012). This appears different from the results here and suggests that there may be considerable differences between the encoding of nearby prefrontal cortical regions. An interesting distinction from the function of other medial prefrontal cortical regions is that vmOFC activity does not control the expression of learned licking behavior (Namboodiri et al., 2019), but regions such as the prelimbic cortex do (Otis et al., 2017; Parent et al., 2015).

Another major unresolved question is how OFC controls behavioral learning rate by interacting with the midbrain dopaminergic neurons signaling RPE (Schultz et al., 1997; Steinberg et al., 2013). OFC has been shown to project heavily to the striatum (Gremel et al., 2016; Groman et al., 2019; Pascoli et al., 2018). This projection may be particularly important for an interaction with the dopaminergic system. Though we have investigated vmOFC→VTA neurons in this and our previous study (Namboodiri et al., 2019), this projection passes through the striatum, and may send collaterals to the striatum. Thus, we cannot rule out the possibility that collaterals in the striatum mediate some of the effects of vmOFC→VTA inhibition in our previous study (Namboodiri et al., 2019). One especially promising circuit mechanism may be via the low threshold spiking interneurons in dorsal striatum, which show similar reduction in reward responses during instrumental learning (Holly et al., 2019). If true, this would raise an interesting possibility that the role of this output and OFC in the control of habitual behavior or behavior insensitive to negative outcomes (Gremel et al., 2016; Groman et al., 2019; Morisot et al., 2019; Pascoli et al., 2018) may be due to its role in controlling learning rate (e.g. learning rate to negative outcomes set too low). OFC could also control behavioral learning rate through its interactions with anterior cingulate cortex (Soltani and Izquierdo, 2019).

Though the circuit mapping experiments performed here cannot ascertain whether the role of mThal input to vmOFC is unique among other inputs, three features of the observed results are worth highlighting. One, inhibition of mThal input to vmOFC changes activity in vmOFC clusters proportional to the amount of adaptation in reward responses (**Fig 6E**). In fact, we showed that 50% of the cluster-wise variance in the effect of mThal inhibition is explained by the variance in adaptation between clusters. Such a robust causal relationship between mThal input and vmOFC output suggests that mThal might indeed be one of main inputs contributing to the reward response adaptation observed in vmOFC. Two, a concern could be that that any strong input with reward activity would control reward response adaptation in vmOFC. An implicit assumption of this concern is that the observed effect is simply a reflection of the strength of reward encoding. The cluster with the strongest reward response in vmOFC is cluster 5 (**Fig 2, 3**). If a strong input to vmOFC contributes to the strongest output (i.e. cluster 5), we would expect a significant change in the activity of cluster 5 due to input inhibition. Yet, this cluster had one of the lowest effects of mThal→vmOFC inhibition (**Fig 6**). Thus, the effect of inhibition is not simply related to the strength of vmOFC activity. In this light, it is unlikely that any strong input would reproduce this effect. Lastly, thalamic outputs are excitatory (Halassa and Sherman, 2019) and exhibit positive reward responses (**Fig 5F**). Yet, inhibition of mThal→vmOFC input increases neuronal response to reward in vmOFC excitatory neurons. These results imply a key role for vmOFC inhibitory interneurons in shaping the responses of vmOFC output neurons. This means that while mThal inputs are integral for the reward response adaptation in vmOFC output neurons, the computation happens within the vmOFC local circuit. Despite these noteworthy features of the mThal input to vmOFC, future work needs to address whether other inputs contribute to learning rate control by vmOFC. For instance, dorsal raphe serotonergic neurons have been shown to control learning rates (Grossman et al., 2020) and may potentially do so through their interactions with OFC.

A potential confound for the response change in vmOFC due to inhibition of mThal→vmOFC input is that the observed vmOFC response changes may be due to a change in behavior resulting from the input inhibition. However, this is unlikely as the inhibition we performed is unilateral and restricted to the field of view under the lens. Indeed, based on tissue scattering and the LED power, it is likely that the inhibition of thalamic axons occurs only within 200 μm or less of tissue under the lens. In the behavior we measured (licking), we observed no effect of inhibition of thalamic axons (**Fig S6C**). Considering that behavioral effects in prefrontal circuits are typically only apparent in strong bilateral inhibition/lesion, it is highly unlikely that the small-scale inhibition in the local circuit under the lens produced any unmeasured behavioral effects that indirectly modulated vmOFC neuronal responses.

In conclusion, we have shown that vmOFC reward responses signal the learning rate for rewards. Whether these results generalize to a broad role for OFC in prioritizing environmental stimuli for learning remains to be tested. Nevertheless, the identification of mThal→vmOFC circuit as one involved in the control of learning rate opens the possibility to study the neural circuit control of parameters for learning, i.e. meta learning.

## Acknowledgments

We thank P. Phillips, J. Berke, D. Ottenheimer, A. Mohebi, K. Ishii, R. Gowrishankar, S. Piantadosi, J. Rodriguez-Romaguera and Z.C. Zhou for comments on the manuscript, and S. Mihalas and all Stuber lab members for helpful discussions. We thank Karl Deisseroth (Stanford University) and the GENIE project at Janelia Research Campus for viral constructs.

## Author Contributions

V.M.K.N., T.G.H., I.T.P., R.S., and M.M.G. performed experiments. V.M.K.N. and M.M.G. performed analyses. V.M.K.N., and G.D.S. designed experiments and wrote the manuscript with input from all authors.

## Data and materials availability

The data used in this study are available from the corresponding author upon request. All the behavioral data were collected using custom MATLAB and Arduino scripts written by VMKN. These are available upon request. The analyses were done in Python using custom codes written by VMKN. These will be uploaded to the Stuber lab Github page (https://github.com/stuberlab), and/or will be available upon request from the corresponding author.

## STAR Methods

### Subjects and Surgery

All experimental procedures were approved by the Institutional Animal Care and Use Committee of the University of North Carolina and University of Washington and accorded with the Guide for the Care and Use of Laboratory Animals (National Institutes of Health). Adult male and female wild type C57BL/6J mice (Jackson Laboratories, 6-8 weeks, 20-30 g) were group housed with littermates and acclimatized to the animal housing facility until surgery. Survival surgeries were stereotaxically performed while maintaining sterility, as described previously (Namboodiri et al., 2019; Resendez et al., 2016). Induction of anesthesia was carried out by using 5% isoflurane mixed with pure oxygen (1 L/min) for roughly thirty seconds to a minute, after which anesthesia was maintained using 0.6-1.5% isoflurane. The surgeon monitored respiratory rate intermittently to ensure appropriate depth of anesthesia. The animals were placed on a heating pad for thermal regulation. Data from animals used in **Fig 1-4**, **Fig S1, Fig S2 and Fig S3A, B** were collected at UNC. For these animals, pre-operative buprenorphine (0.1 mg/kg in saline, Buprenex) treatment was given for analgesia. Eyes were kept moist using an eye ointment (Akorn). 2% lidocaine gel (topical) or 1mg/kg lidocaine solution was applied or injected onto the scalp prior to incision. Details of viral injection, lens and optic fiber implantation are provided below. A custom-made stainless-steel ring (5 mm ID, 11 mm OD, 2-3 mm height) was implanted on the skull for headfixation and stabilized with skullscrews as well as dental cement. Animals either received acetaminophen (Tylenol, 1 mg/mL in water) in their drinking water for 3 days, or 5 mg/kg carpofen 30 min prior to termination of surgery for post-operative analgesia. Animals were given at least 21 days (and often, many more) with ad libitum access to food and water to recover from surgery. Following recovery, animals used for behavioral studies were water deprived to reach 85-90% of their pre-deprivation weight and maintained in a state of water deprivation for the duration of the behavioral experiments. Animals were weighed and handled daily to monitor their health. The amount of water given daily was between 0.6-1.2 mL and was varied for each animal based on the daily weight. A total of 38 mice were included in this study: 5 OFC-CaMKII imaging (0 female, **Fig 1-3**), 7 OFC-VTA imaging (0 female, **Fig S2, 3**), 5 mThal→vmOFC axon imaging (3 female, **Fig 5, Fig S5**), 7 mThal→vmOFC inhibition or control inhibition during vmOFC imaging (3 female, **Fig 6**, **Fig S6**), 10 anatomical tracing (8 female, **Fig 5B**), and 4 pupil diameter tracking (0 female, **Fig S4**).

### Head-fixed behavior

Trace conditioning was done exactly as before (Namboodiri et al., 2019). A brief outline of these methods is summarized here. Water deprived mice were first trained to lick for random unpredictable sucrose (10-12.5%, ∼2.5 µL) deliveries in a conditioning chamber. Mice received one of two possible auditory tones (3 kHz pulsing tone or 12 kHz constant tone, 75-80 dB) that lasted for 2 seconds. A second after the cues turned off, the mice received a sucrose reward following one of the tones (designated CS+), whereas the other tone resulted in no reward (designated CS−). The identity of the tones was counterbalanced across mice in all experiments. The cues were presented in a pseudorandom order and in equal proportion until a total of 100 cue presentations (trials) were completed. The intertrial interval between two consecutive presentations of the cues was drawn from a truncated exponential distribution with mean of 30 s and a maximum of 90 s, with an additional 6 s constant delay. Early in learning (**Fig 1**) was defined as the first session of conditioning. Late in learning (**Fig 1**) was defined as the day that the area under a receiver operating characteristic curve (auROC) of lick rates to CS+ versus CS− remained high and stable (auROC larger than 0.7 on at least 2 consecutive sessions or larger than 0.85). Two contingency degradation experiments were performed as described previously (Namboodiri et al., 2019), with reward probability reduced 50% in one (**Fig 1,2**) and background unpredictable rewards introduced in the intertrial interval in the other, with reward probability set to 100% (**Fig S2**, **Fig 3**). Exact parameters for the rates of unpredicted rewards were described previously (Namboodiri et al., 2019). For the sucrose and quinine experiment (**Fig 3**), drops of sucrose (10-12.5%, ∼2.5 µL) or quinine hydrochloride dihydrate (1.5-2.5 μM, ∼2.5 µL) were randomly delivered at a 3:1 ratio (60 sucrose drops and 20 quinine drops). The interdrop interval was a minimum of 13-18 s and a maximum of 23-28 s. These were chosen to maintain a sufficient interval between consecutive drops so as to prevent any bleed-through of GCaMP fluorescence. Even though the hazard rate is not flat for these intervals, the animals did not show any behavioral evidence of temporal expectation of the delivery times.

### 2-photon microscopy

The methods were similar to those published previously (Namboodiri et al., 2019). We used a calcium indicator (GCaMP6s) to image calcium changes using 2-photon microscopy. The injection coordinates and volumes for virus as well as the coordinates for implanting a gradient refractive index (GRIN) lens were as published previously (Namboodiri et al., 2019). For mThal axon imaging (**Fig 5**) or inhibition (**Fig 6**), we injected 400-500 nL of AAVDJ-CaMKIIα-GCaMP6s (at an effective titer of ∼1-2×10^12^ infectious units per mL, UNC Vector Core) or AAV5-CaMKII-eNpHR3.0-mCherry (∼4×10^12^ infectious units per mL, UNC Vector Core) unilaterally in mThal (−1.3 AP, 0.5 ML, −3.5 DV from bregma). We used a resonant scanner (30 Hz frame rate acquisition, Olympus Fluoview FVMPE-RS) and performed an online averaging of 6 times to get an effective frame rate of 5 Hz, to minimize the size of recorded files as we had negligible motion artifacts. A GaAsP-PMT with adjustable voltage, gain and offset was used, along with a green filter cube. We used a long working distance 20x air objective, specifically optimized for infrared wavelengths (Olympus, LCPLN20XIR, 0.45 NA, 8.3 mm WD). We imaged either at 955 or 920 nm using a Ti-sapphire laser (SpectraPhysics, ∼100 fs pulse width) with automated alignment. The animals were placed on a 3-axis rotating stage (Thorlabs, TTR001/M) to precisely align the surface of the GRIN lens to be perpendicular to the light path, such that the entire circumference of the lens is crisply in focus (within 1-2 µm). The imaging acquisition was triggered by a custom Arduino code right before the start of a behavioral session, and a TTL output of every frame was sent as an input to the Arduino. The imaging acquisition was triggered off at the end of the behavioral session (∼ one hour). The data in **Fig 1, 2** (OFC-CaMKII) and **Fig S2** (OFC→VTA) were re-analyzed based on data from a previous publication (Namboodiri et al., 2019). In every mouse, one z-plane was imaged throughout acquisition so that the same cells could be tracked through learning (Namboodiri et al., 2019). After mice were trained, other z-planes were also imaged (one per session) to get a measure of the total functional heterogeneity in the network. A total of 2-6 z-planes per mouse were imaged in the OFC- CaMKII group, whereas 1-3 z-planes were imaged in the OFC-VTA group. Thus, responses early in learning were from the plane tracked throughout learning, whereas the responses late in learning were from all imaged planes. The sucrose quinine session was run for most imaging planes at the end of the conditioning experiments (Namboodiri et al., 2019). The data in **Fig 3** were collected from the same animals as in **Fig 1, 2**, but have never been published previously.

### Imaging data analysis

Preprocessing (motion correction, manual ROI detection, signal extraction, neuropil correction) using SIMA (Kaifosh et al., 2014) for vmOFC cell body imaging was as described previously (Namboodiri et al., 2019). For axon imaging, we used the same approach as for cell bodies for motion correction. For detection of axonal ROIs, we employed a manual hand-drawing method but with different criteria: 1) we drew ROIs only around parts of an axon that showed no resolvable overlap with other fluorescent regions, thereby making the ROIs small, 2) we drew ROIs along a single axon (at least those that can reliably be tracked along the imaging focal plane) only once (more on this below), 3) we drew ROIs around regions of an axon that are definitely within the imaging plane (these are often bright and sharply in focus), 4) we drew ROIs that were at least 10 pixels or so, in order to minimize noise. An example plane with manually annotated ROIs is shown in **Fig S5**. Despite these precautions, it is impossible to know whether different ROIs are from the same underlying axon, as axons move in and out of the imaging plane and are highly branched. Thus, we do not make any claims about the individual ROIs shown in **Fig 5F** representing individual axons (though the example traces in **Fig 5E**, representing the axonal ROIs with the largest skew in activity, show distinct patterns). A common approach to identify ROIs that are putatively from the same axon is to remove segments that show high correlations. However, this threshold depends crucially on the signal to noise ratio of the recording. If the signal to noise ratio is low, segments of the same axon will show low correlations due to the noise dominating the fluorescence. Thus, the decision to set the threshold often becomes subjective, especially considering variability in signal to noise ratio between animals resulting from variability in GCaMP expression. We avoid this problem by not claiming that different ROIs are necessarily from different axons. Instead, we quantify both a population mean of all ROIs and separately perform a clustering analysis of ROIs based on their response profiles to identify heterogeneous response profiles (more on this below). We did not perform any neuropil correction for axon recordings as the neuropil is primarily due to signals of interest (i.e. from axons). Due to this reason, our earlier approach of ensuring no overlap with other axons is important to get isolation of signals.

The clustering analysis for vmOFC neurons was performed as previously published (Namboodiri et al., 2019). Importantly, all clustering was performed based on the peristimulus time histograms (PSTHs) around CS+ and CS− in the session after learning with 100% reward probability. So, any neuron that has an assigned cluster identity was recorded under the 100% contingency late in learning. Once a cluster ID was assigned to a neuron, the same cluster ID was used for that same neuron in all other sessions. This was possible because 2-photon imaging allowed longitudinal tracking of the exact same set of neurons across many days and tasks (Namboodiri et al., 2019).

The clustering of axon ROIs (**Fig S5**) used the same approach but used PSTHs around sucrose alone (**Fig S6G**), or around both sucrose and quinine (**Fig S5H**). As we did not longitudinally track axons across these sessions, we performed clustering separately for these two sessions. Nevertheless, qualitative correspondence between the sucrose responses of some clusters can be seen by comparing **Fig S5G** and **H**. The benefit of performing cluster analysis for axons is that different identified clusters are almost certain to be from different underlying axons, as the clustering is done on average responses across trials, thereby reducing noise. Thus, we do not assess axon-by-axon heterogeneity as we cannot reliably identify individual axons. We can nevertheless assess heterogeneity of information encoding in the axonal population by interpreting cluster-wise differences in response profiles. This is philosophically similar to our approach with clustering of vmOFC neurons. We only interpret average results across all neurons within a cluster, thereby treating a cluster as a unit of information representation. The one big difference between the identification of cell body and axon clusters is that unlike in the cell body case, we cannot assess prevalence of each cluster among the axonal population. This is because the ROIs making up any cluster might potentially be overlapping and represent the same axon.

In **Fig 2**, we quantified reward responses as the coefficients of a General Linear Model (GLM) fit to reward delivery (Namboodiri et al., 2019). Importantly, this GLM approach was applied to the deconvolved calcium fluorescence to remove fluorescence changes purely due to the dynamics of GCaMP6s. We employed a GLM approach primarily to separate lick related responses and reward prediction or receipt responses, as both licking and rewards generally produced positive responses (Namboodiri et al., 2019). We did not employ a similar GLM approach for analyzing the sucrose and quinine responses in **Fig 3** as these responses were evidently dissociated from licking responses. This is because licking was high for sucrose alone, high for sucrose in the sucrose-quinine session and low for quinine in the sucrose-quinine session; yet, the responses were generally high, low and high respectively, for these conditions.

To obtain the average population slope in **Fig 1L**, we first fitted a best-fit linear regression to the trial-by-trial variability in OFC reward response (separately in rewarded and unrewarded trials) with the licking update on the next trial for each neuron. We then averaged these slopes across all recorded neurons to obtain the average population slope separately for the positive RPE case (i.e. rewarded trials in the 50% reward probability session) and the negative RPE case (i.e. unrewarded trials in the 50% reward probability session). To obtain the visualization plot shown in **Fig 1K**, we first z-scored both the trial-by-trial variability of a neuron and the licking update (i.e. both axes) for each neuron and pooled these data for all neurons. We then binned the data along the activity z-score and calculated the corresponding mean lick update for that bin (using numpy.digitize with right=False, i.e., a datapoint is counted in a given bin if it is greater than or equal to the lower edge of the bin and less than the higher edge). The plot shows the lower edges of the bins, with z < - 3 not shown due to low sampling. The z-scoring was performed prior to pooling to avoid confounding within-neuron and between-neuron variability. The actual quantification was done on the raw slopes instead of the z-scored data to avoid equating neurons with considerable difference in the trial-by-trial variability of reward responses.

### Optogenetics during imaging

These animals received an injection of AAVDJ-CaMKIIα-GCaMP6s in vmOFC and AAV5-CaMKIIα- eNpHR3.0-mCherry (experimental) or AAV5-CaMKIIα-mCherry (control) in mThal. All animals showed significant expression of the opsin in mThal. Light was delivered for optogenetic inhibition to the full field of imaging by an LED kit with a peak wavelength of 615 nm (FV30SP-LED615, Olympus) (Otis et al., 2019). The frames containing light artifacts due to LED illumination were masked out for analysis (shown as white bars in **Fig 6B, C**). The same preprocessing pipeline as before (including motion correction, signal extraction and neuropil correction) was employed on this masked data. These animals were first trained to lick in response to random sucrose deliveries. We then performed optogenetic inhibition during sucrose consumption for multiple imaging planes. A random half of the trials received inhibition. The effect of the LED was calculated by comparing sucrose fluorescence on the trials with and without LED. The animals were subsequently trained on the trace conditioning paradigm (no imaging). Once anticipatory licking was high and stable, we imaged the same neurons that were imaged earlier to obtain PSTHs around CS+ and CS−. These PSTHs were used to classify neurons into the clusters identified using the much larger population of neurons in **Fig 2**. The classification was done using a linear support vector classifier (Scikitlearn), as was used previously for classifying OFC→VTA neurons (Namboodiri et al., 2019). We then performed inhibition of mThal axons while imaging from vmOFC neurons during the sucrose and quinine session (**Fig 6C, D**). Here, to obtain a sufficient number of trials with and without inhibition, we first performed recordings without inhibition (80 trials), followed by with inhibition (80 trials). The effect of LED was calculated by comparing sucrose or quinine fluorescence with and without LED.

### Retrograde tracing, histology and microscopy

400 nL of rAAV2retro-hSyn-eYFP (∼2×10^12^ infectious units/mL) or CTB-488 were stereotaxically injected at roughly 2.5 AP, 0.5 ML and 2.3-2.5 DV from bregma using the surgical methods described above. 3-5 weeks after surgery, animals were euthanized with an overdose of pentobarbital (∼390 mg/kg, Somnasol, Covetrus EU-HS-045-100-0), and transcardially perfused with 4% paraformaldehyde (PFA, Sigma-Aldrich, #158127). Perfused brains were incubated in 4% PFA overnight and moved to a 30% sucrose solution (Sigma-Aldrich, #S0389) for ∼2 days prior to cryosectioning. 40 µm thick sections were used in tracing experiments. For retroAAV2 thalamic labeling (**Fig 5B, Fig S5**), eYFP signal was enhanced and stabilized using a chicken anti-GFP antibody (Aves Lab, #GFP-1020, 1:500 dilution), paired with a donkey anti-chicken secondary (Jackson Immunoresearch, #703-545-155, 1:1000 dilution). GFP and eYFP have highly similar protein sequences, which allows the use of a GFP antibody for immunostaining. Brain sections were imaged using a 20x air objective on a confocal microscope (Olympus Fluoview FV3000). Resulting image tiles were stitched, and Z-stacks were taken at ∼1 µm intervals and averaged across slices yielding a maximum intensity projection image. Brain atlas outlines (https://mouse.brain-map.org/static/atlas) were overlaid onto each image to allow assignment of thalamic subregions, in which labeled cells were counted using ImageJ (https://imagej.net/Fiji). Percentage of total thalamic cells labeled (**Fig 5B**) was quantified as the number of cells per region divided by the sum total of all counted eYFP+ or CTB+ cells. The intermediodorsal nucleus was counted as part of the mediodorsal region.

### Pupil measurements

Pupil area measurements were performed on an independent cohort of animals that went through the complete head-fixed behavior paradigm. Analysis was performed for the unpredicted sucrose without quinine condition, the early in learning condition, the 50% reward probability and the unpredicted sucrose and quinine delivery conditions. Pupil recordings were performed using a monochrome USB 2.0 CMOS Camera (ThorLabs DCC1545M) at 5 Hz. A triggered red LED flash in the inter-trial interval was used to align behavior and camera recordings. This flash occurred outside of the analyzed window for all recordings. To align the behavior and pupil recordings, LED flashes were detected using ImageJ and MATLAB to identify large fluctuations in pixel intensity that corresponded to LED onset from an ROI that contained the LED. The LED onset timestamps for the video recordings were aligned to the LED trigger timestamp in the behavior recordings on a trial by trial basis to account for dropped frames across the session, though very few frames were dropped overall. After the video and behavior recordings were aligned, we extracted data from the interval spanning 3 seconds before cue presentation to 3 seconds after reward delivery for the early in learning and 50% reward probability conditions, and 3 seconds before and after the first lick following uncued fluid delivery for the unpredicted sucrose without quinine and unpredicted sucrose with quinine sessions. We preprocessed the data with two runs of the CLAHE ImageJ plug-in (Zuiderveld, 1994) with default parameters to enhance local contrast for pupil discrimination. After preprocessing, we performed an average intensity grouped z-projection of 5 frame bins to reduce the data to 1 Hz for manual annotation of the pupil. We then drew an ellipse bounding the pupil for each resulting frame and extracted the area of the ellipse for each frame. For sessions with multiple trial types, we did not attach trial identity data to each trial until all pupil measurements were complete.

## Supplementary Information

### Supplementary Note 1

It may be worth noting that in addition to the salience related to aversion, quinine may also have been salient in the sucrose-quinine experiment for two additional reasons. One, the mice in our task had considerable experience receiving sucrose under the two-photon microscope. However, quinine was a novel stimulus in the sucrose-quinine session. Novelty typically increases salience. Two, the presence of quinine reduces the context-reward association such that a high frequency of quinine could make licking in the context aversive, thereby making an estimation of quinine frequency more salient for deciding whether to lick. All three of these reasons cause sucrose to be relatively less salient for learning in the presence of quinine.

### Supplementary Note 2

#### Discussion of all major findings in the study and their support or falsification of various models/explanations

We consider several possible alternative models here. In general, model comparisons require two steps. The first step is to rule out models that are not consistent with the data. The second step is to quantitatively compare the remaining models that fit the data based on parsimony (e.g. number of free parameters, information criteria, cross validation etc.). <u>Here, we only undertake the first step, as we show that encoding of most variables commonly assumed in reinforcement learning is inconsistent with the full set of vmOFC reward response observations.</u> Of course, much more complex alternatives that incorporate mixtures of many variables can possibly fit all observations. Nevertheless, we believe that the strongest support for learning rate control comes from the data shown in **Fig 1**. These data can only be accounted for by models that at least partly assume learning rate control. Learning rate control is also consistent with all the other observations, as we explain below. Thus, we believe that the most parsimonious account of our data is a monotonic encoding of learning rate. We first provide an account of the common alternative variables assumed in reinforcement learning before exhaustively discussing all possibilities.

##### Common alternative reinforcement learning variables

###### RPE

Since RPE is a linear combination of reward receipt and reward prediction, the results in **Fig S2** rule out RPE. We summarize these results here. Due to temporal uncertainty, RPE for a delayed reward does not become zero even after extensive training. Under similar delays as employed here, prior findings show that midbrain dopaminergic neurons exhibit significant positive responses to a predicted reward even after extensive training (Coddington and Dudman, 2018; Cohen et al., 2012; Engelhard et al., 2019; Kobayashi and Schultz, 2008; Lee et al., 2020). However, we observed negative responses in multiple clusters (especially 1 and 3) to reward after learning **(Fig 2D, E)**. Further, reward omission should produce a negative RPE, and unpredicted reward delivery should produce a positive RPE independent of whether the session contains an unpredicted aversive stimulus. These predictions are violated by the responses of many clusters (**Fig S2** and **Fig 3D-G**).

###### Absolute magnitude of RPE

The Pearce-Hall model of surprise proposes that learning rate is controlled by the absolute magnitude of RPE (Pearce and Hall, 1980). Based on this model, the absolute magnitude of the RPE for unpredicted sucrose should not reduce merely due to the presence of quinine in the context. It should also be high and positive for both unpredicted sucrose and quinine. The absolute magnitude of the responses of midbrain dopamine neurons, a readout of RPE, match these predictions. For instance, a previous study evaluated the response of midbrain dopaminergic neurons to both rewarding and aversive stimuli while varying the general probability of reward (Matsumoto et al., 2016). They found that reward responses of dopamine neurons were high despite the presence of aversive stimuli. They also found that while the response to the aversive stimulus (a salient air puff) was positive or negative dependent on the general reward probability, the magnitude of response was high in all cases. Thus, a signaling of the absolute magnitude of the RPE should produce high and positive responses to both unpredicted sucrose and quinine. Instead, we observed that unpredicted sucrose response reduces in a context containing unpredicted quinine (**Fig 3D-G**). Thus, even though signaling of the absolute magnitude of RPE would be consistent with the data in **Fig 1**, vmOFC reward responses do not correspond to the absolute magnitude of RPE.

A previous study observed a reduction in dopamine responses to reward by the induction of acute stress or fearful states (Zhong et al., 2017). Hence, it may be possible to argue that the quinine context is aversive or stressful and hence, would result in a reduction of the absolute magnitude of RPE to sucrose. However, this argument is highly unlikely to be true. This is because we introduced quinine in only one out of four liquid deliveries. The remaining are all sucrose. Such a low fraction of quinine presentations is highly unlikely to induce an acute stressful state. Indeed, the animals remain highly motivated to lick and consume the sucrose under this condition (**Fig 3B**). Further, (Matsumoto et al., 2016) observed strongly positive reward responses in dopamine neurons even when <20% of outcomes were rewarding. In comparison, 75% of the outcomes in the quinine context were rewarding in our experiment. Overall, absolute magnitude of RPE is not consistent with these observations from vmOFC neurons.

###### Subjective value of received reward

Activity in vmOFC is not consistent with a signaling of the subjective value of a received reward in these experiments (Amarante and Laubach, 2020; Padoa-Schioppa and Assad, 2006). This is because the subjective value of the same received reward should be equal regardless of whether the reward was predicted, and further the subjective value of the receipt of a predicted reward should be larger than the subjective value of the omission of a predicted reward. Lastly, the subjective value of an aversive stimulus should be lower than that of a rewarding stimulus. All these predictions are violated (**Fig S2, Fig 3**).

###### Relative value of received reward

It has previously been observed that orbitofrontal cortical neurons represent the relative value of a reward in comparison to other available options (Tremblay and Schultz, 1999) (though see (Amarante and Laubach, 2020; Padoa-Schioppa and Assad, 2008)). Such a relative value signal has also been clearly demonstrated in other brain regions (Louie et al., 2011; Ottenheimer et al., 2018). If vmOFC reward responses reflect relative value, when a reward is presented along with a highly non-preferred option (quinine), the response to the reward should *increase*, as the relative value of sucrose in relation to quinine is higher than when sucrose is presented alone. Instead, we observed that the response to unpredicted sucrose reduces in a context containing unpredicted quinine (**Fig 3D-G**). Further, we also observed that unpredicted quinine produces a larger response than unpredicted sucrose (**Fig 3D-G**). Thus, these results rule out relative value encoding.

###### State value

vmOFC reward responses are also not consistent with the expected future value or the state value. Expected future value is the predicted discounted sum of all future rewards given the current state. In other words, the expected future value after a reward is the expected discounted value of all *future* rewards given that a reward was just received. Prior to learning the cue-reward association, the receipt of a reward should produce a positive expected future value since it raises the possibility that there will be more future rewards. After learning the cue-reward association, the animals should have a high expected future value following cue presentation, but low expected future value once the reward is received. This is because once the predicted reward is obtained, the intertrial interval contains no reward; only the next cue presentation predicts a reward. These predictions are qualitatively supported by vmOFC responses as their reward responses generally reduce after learning (**Fig 2**). Nevertheless, the expected future value after a reward should be even lower when the reward probability is reduced to 50%, as the overall reward rate is lower and the animals have to wait longer to receive the next reward. However, we observed that the responses on rewarded trials of clusters 1 and 3 are higher in the 50% probability session compared to the 100% probability session (**Fig S2**). Additionally, expected future value should also be equal to both the unpredicted sucrose and the unpredicted quinine in the sucrose and quinine session since both events predict a similar future value (as neither predicts the upcoming sequence of stimuli). This was also not observed (**Fig 3D, E**).

###### Behavioral confounds

The observed results also rule out simple behavioral confounds. Specifically, the sucrose-quinine experiments rule out confounds due to variability in licking or a lingering taste of quinine. Variability in licking to sucrose and quinine cannot explain the reduction in unpredicted sucrose responses in a context containing quinine. This is because the consumption lick rate is high for unpredicted sucrose without quinine, high for unpredicted sucrose in a context containing quinine, and low for unpredicted quinine. However, the responses are high for unpredicted sucrose without quinine, low for unpredicted sucrose in a context containing quinine, and high for unpredicted quinine. A lingering taste of a previous quinine drop after tasting sucrose also fails to explain the results we observed. This is because quinine produced a positive response much like what sucrose produced when delivered alone (**Fig 3D, E**). If the taste of sucrose and quinine interact, it should produce a sucrose response similar to that of the quinine response. Instead, we observed that the sucrose response diverged from the quinine response, a finding that cannot be explained by simple history effects of taste. Lastly, vmOFC reward responses do not merely reflect an efference copy of an arousal signal controlling pupil diameter. We found that the mean pupil diameter responses to sucrose early in learning and after 50% reward probability reduction do not correlate with the corresponding vmOFC reward responses (**Fig S4**). Nevertheless, consistent with a reduction in the relative salience of sucrose in a context containing quinine, we observed a highly dampened pupil dilation for sucrose in this context (**Fig S4**). These observations show that a simple efference copy of an arousal signal is not sufficient to explain vmOFC reward responses.

##### Extensive consideration of all major findings in the manuscript in relation to many models

We first list the various possible models for the data and then list the various major findings in the manuscript. We consider each model against each observation later.

##### Possible models/explanations for the data

1. Learning rate through relative salience
2. Learning rate through Pearce Hall salience, i.e. absolute magnitude of previous trial’s RPE. This model is the closest alternative to the data, as it involves the control of learning rate, the central claim in the manuscript. However, Pearce Hall salience is explicitly formalized as the absolute magnitude of the previous trial’s RPE. We will consider whether this explicit formalism provides a parsimonious explanation below.
3. Reward Prediction Error
4. Reward value
5. Expected Future Value: this is the discounted sum of future rewards given that the reward was just received (or omitted).
6. Updated state value after current outcome: this is the expected future value of the state immediately before reward delivery plus learning rate multiplied by the RPE at the time of reward. In other words, this is the state value immediately prior to the outcome after having observed the outcome.
7. Relative value
8. Variability in licking
9. Variability in pupil diameter or arousal
10. Reward value plus lingering taste
11. Reward value plus satiation
12. Identity prediction error: this is a prediction error about the identity of the outcome without a consideration for its value.

<u>Major findings in our study</u> and whether they support or falsify each model

*Please note that in general when additional assumptions are required for a model, we have assumed the conditions that produce predictions most closely resembling the observations. In other words, alternative models are ruled out in a conservative manner*.

1. <u>There is a positive correlation between vmOFC reward responses (rewarded trials) on one trial and the change in anticipatory licking on the next CS+ trial from the current CS+ trial (**Fig 1J-L**).</u> A mathematical model of the role of learning rate in reward prediction updating can be described as Reward prediction on trial n+1 of CS+ cue = reward prediction on trial n of CS+ cue plus (learning rate on trial n) multiplied by (RPE on trial n) (shown in **Fig 1A**) If vmOFC activity is controlling learning rate, lick rate should change between two consecutive CS+ trials roughly proportional to vmOFC reward activity on that trial multiplied by the RPE on that trial. This assumes a monotonic relationship between reward prediction on a trial and lick rate on that trial. On rewarded trials during 50% probability sessions, RPE should be positive. Hence, trial-by-trial changes in learning rate (hypothesized to be vmOFC reward time activity on the trial) should produce a *positively correlated* change in lick count update. In other words, lick count change between two consecutive CS+ trials should *positively* depend on vmOFC reward response on that trial (representing the learning rate). This rationale is explained in **Fig 1**. Thus, this observation is consistent with the learning rate models (relative salience and absolute magnitude of RPE). It is also consistent with the functional consequence of every model but the variability in licking and identity prediction error models.
2. <u>There is a negative correlation between vmOFC reward expectation response (unrewarded trials) on one trial in the 50% condition and the change in anticipatory licking on the next trial from the current trial (Fig 1J-L).</u> On unrewarded trials during 50% probability sessions, RPE should be negative. Hence, lick count change between two consecutive CS+ trials should *negatively* depend on OFC reward omission response on that trial under the learning rate models (relative salience and absolute value of RPE). Thus, both these models are supported by this observation. Every other model predicts either no dependence or a positive dependence (e.g. **Fig 1C**).
3. <u>vmOFC responses are positive to reward across all clusters prior to learning (Fig 2D, E)</u>. All models are consistent with this observation.
4. <u>vmOFC responses are negative to reward or considerably reduced across many clusters post learning in the 100% reward probability condition (Fig 2D, E).</u> Learning rate through relative salience is consistent with this observation since a predicted reward is less salient for learning. We assume that a monotonic function of the reward response will control learning rate, such that negative reward responses result in near-zero learning rate. Expected future value is also qualitatively consistent with this observation since the animals would have learned that there is no reward available in the intertrial interval. None of the other models are consistent with this observation. RPE for a delayed reward should be significantly positive even after learning and so should absolute value of RPE and identity prediction error. Thus, the activity at least in clusters 1 and 3 are not consistent with these variables. Reward value and relative value should be the same as in the session early in learning. Updated state value should have stabilized at its peak value and should be highly positive. The animals lick at a high rate to consume the reward delivered early and late in learning and thus, licking cannot explain a drastic reduction in reward response. Pupil diameter is high for reward both before and after learning. Satiation level should be the same early and late in learning since the animals receive the same number of rewards.
5. <u>vmOFC responses are positive to reward omission (**Fig S2A, B**).</u> Learning rate through relative salience or absolute magnitude of RPE are supported by this observation as negative RPEs typically cause larger learning rates than positive RPEs (Frank et al., 2009; Galea et al., 2015; Gershman, 2015; Kojima et al., 1996). Expected future value should be high since the animals would not know whether the reward is omitted or not for some time after reward time due to the temporal uncertainty in subjective reward timing. Updated state value should be positive for the same reason as the expected future value. Animals typically lick at a high rate for some time after reward delay due to temporal uncertainty. Pupil diameter is positive after reward omission. Hence, models 1, 2, 5, 6, 8 and 9 are supported by this observation. The remaining models are falsified. RPE and identity prediction error should be negative. Reward value and relative value should be zero without receiving reward.
6. <u>vmOFC responses are lower for reward receipt trials than reward omission trials for many clusters (**Fig S2A, B**).</u> Learning rate should be lower on positive RPE trials compared to negative RPE trials. Absolute magnitude of RPE could potentially be lower for rewarded compared to unrewarded trials if the reward prediction probability is higher than 50%. Expected future value is low once reward is received, since no more rewards are expected until next cue. Hence, all the above models are potentially supported by this observation. All remaining models are falsified. RPE or identity prediction error should be more positive for rewarded compared to unrewarded trials. All variants of reward value should be higher when reward is received. Updated state value should be higher after reward receipt than after omission. Pupil diameter is lower for reward omission compared to reward receipt.
7. <u>vmOFC responses are positive for unpredicted rewards early and late in learning (Late: Fig 3F; Early: Fig 2D, E, Fig S3).</u> All models are supported by this observation.
8. <u>vmOFC responses are lower for unpredicted sucrose in a context containing quinine (**Fig 3**; compare to early in learning in Fig 3D, E; compare to unpredicted sucrose without quinine late in learning in Fig 3F; population responses for each cluster can also be compared to the unpredicted responses in **Fig S3B**).</u> Learning rate through relative salience is supported by this observation as sucrose is less salient for learning in this condition. Expected future value should be lower in the sucrose and quinine session since the reward rate is lower in this session. All other models are falsified by this observation. RPE should be equal for unpredicted sucrose in a context with or without unpredicted quinine. All variants of reward value should be high. Indeed, relative value should be even higher when sucrose is delivered in a context containing quinine. Identity prediction error should be higher in this condition. When all liquids are predicted to be sucrose, identity prediction error should be zero. Any violation of this expectation should produce a positive prediction error. Updated state value should be high since there is no prediction, but reward value is the same. Interestingly, animals licked to consume sucrose at a higher rate in the sucrose and quinine session compared to the session early in learning (data not shown). Thus, variability in licking cannot explain these results either.
9. <u>vmOFC responses are positive for unpredicted quinine (Fig 3D, E).</u> Quinine is salient and hence, this observation is consistent with the relative salience model. It is also consistent with the absolute magnitude of RPE and an identity prediction error. Every other model is falsified by this observation. All variants of reward value should be positive. RPE should in principle be negative, even though positive responses of dopamine neurons to aversive stimuli have been observed in high reward contexts (likely reflecting salience and not RPE) (Matsumoto et al., 2016). Expected future value should be equal for both unpredicted quinine and unpredicted sucrose in the sucrose and quinine session. Updated state value should be low since quinine is aversive. Animals lick at a low rate when they receive quinine (possibly reflecting the uncertainty in the identity of liquid delivered).
10. <u>vmOFC responses for sucrose are higher on rewarded trials in the 50% reward probability session than on rewarded 100% trials for all clusters with negative reward responses (**Fig S2D**).</u> Variability in the environment should increase learning rate. So, this supports the learning rate by relative salience or absolute magnitude of RPE model. RPE should also be higher in this session as the reward prediction is lower. Reward value plus satiation could in principle be supported by this observation as the animals receive fewer rewards and hence, would be comparatively less sated. Every other model is falsified by this observation. Expected future value should be even lower after reward in the 50% session compared to the 100% session. Reward value and relative value should be the same. Updated state value should be lower. Animals lick to consume received reward at the same rate in these sessions (Namboodiri et al., 2019).
11. <u>vmOFC responses are higher for rewarded background sessions (i.e. sessions containing unpredicted rewards during the inter-trial interval) compared to 100% trials for all clusters with negative reward responses (**Fig S2E**).</u> Variability in the environment should increase learning rate. So, this supports the learning rate by relative salience or absolute magnitude of RPE model. Expected future value should be higher after reward when unpredicted rewards are available in the intertrial interval. Updated state value should also be higher for the same reason. Animals typically lick a lot more in the background session (though consumption lick rate is the same (Namboodiri et al., 2019)) and hence, it is possible that the licking model is supported by this observation. A potential lingering taste of previously received sucrose rewards in the intertrial interval could also increase the reward value of the predicted sucrose reward. Thus, all these models are supported by this observation. Every other model is falsified. RPE, reward value, relative reward value, identity prediction error should be the same in both sessions.
12. <u>vmOFC responses are positive for sucrose in the first few trials of the first sucrose and quinine session prior to experiencing quinine for the first time (Fig 3G).</u> All models are supported by this observation.
13. <u>Inhibition of OFC-VTA reward time responses suppresses trial-by-trial updating of anticipatory licking based on reward history. This demonstrates a causal role for OFC activity in controlling behavioral learning (Fig 8B-E in (Namboodiri et al., 2019)).</u> This observation is consistent with vmOFC representing learning rate, RPE, reward value and its variants and updated state value, as all these variables could contribute to trial-by-trial learning. This observation violates expected future value, identity prediction error and the variability in licking models as these variables should not directly contribute to trial-by-trial learning.
14. <u>Pupil diameter during reward receipt or omission does not correlate with vmOFC reward responses, as they are high for reward early in learning, high for reward receipt in the 50% reward probability session and low for reward omission in the 50% reward probability session (**Fig S4**).</u> This is not included in the table below as this finding is about pupil diameter variation and not vmOFC reward response.

In the table below, we summarize whether each of the 12 possible models/explanations can explain the 13 major findings related to vmOFC reward responses. We show that the only model that explains every observation is the one we proposed. Thus, we believe that all the major findings related to vmOFC activity (#1-#14 above) strongly support vmOFC governing the learning rate.

|  | Find-<br>ing<br>1 | Find-<br>ing<br>2 | Find-<br>ing<br>3 | Find-<br>ing<br>4 | Find-<br>ing<br>5 | Find-<br>ing<br>6 | Find-<br>ing<br>7 | Find-<br>ing<br>8 | Find-<br>ing<br>9 | Find-<br>ing<br>10 | Find-<br>ing<br>11 | Find-<br>ing<br>12 | Find-<br>ing<br>13 |
| --- | --- | --- | --- | --- | --- | --- | --- | --- | --- | --- | --- | --- | --- |
| Model<br>1 | ✓ | ✓ | ✓ | ✓ | ✓ | ✓ | ✓ | ✓ | ✓ | ✓ | ✓ | ✓ | ✓ |
| Model<br>2 | ✓ | ✓ | ✓ | ✗ | ✓ | ✓ | ✓ | ✗ | ✓ | ✓ | ✗ | ✓ | ✓ |
| Model<br>3 | ✓ | ✗ | ✓ | ✗ | ✗ | ✗ | ✓ | ✗ | ✗ | ✓ | ✗ | ✓ | ✓ |
| Model<br>4 | ✓ | ✗ | ✓ | ✗ | ✗ | ✗ | ✓ | ✗ | ✗ | ✗ | ✗ | ✓ | ✓ |
| Model<br>5 | ✓ | ✗ | ✓ | ✓ | ✓ | ✓ | ✓ | ✓ | ✗ | ✗ | ✓ | ✓ | ✗ |
| Model<br>6 | ✓ | ✗ | ✓ | ✗ | ✓ | ✗ | ✓ | ✗ | ✗ | ✗ | ✓ | ✓ | ✓ |
| Model<br>7 | ✓ | ✗ | ✓ | ✗ | ✗ | ✗ | ✓ | ✗ | ✗ | ✗ | ✗ | ✓ | ✓ |
| Model<br>8 | ✗ | ✗ | ✓ | ✗ | ✓ | ✗ | ✓ | ✗ | ✗ | ✗ | ✓ | ✓ | ✗ |
| Model<br>9 | n/a | n/a | ✓ | ✗ | ✓ | ✗ | ✓ | ✓ | ✓ | n/a | n/a | n/a | n/a |
| Model<br>10 | ✓ | ✗ | ✓ | ✗ | ✗ | ✗ | ✓ | ✗ | ✗ | ✗ | ✓ | ✓ | ✓ |
| Model<br>11 | ✓ | ✗ | ✓ | ✗ | ✗ | ✗ | ✓ | ✗ | ✗ | ✓ | ✗ | ✓ | ✓ |
| Model<br>12 | ✗ | ✗ | ✓ | ✗ | ✗ | ✗ | ✓ | ✗ | ✓ | ✗ | ✗ | ✓ | ✗ |

---

**Fig S1 (related to Fig 1):**
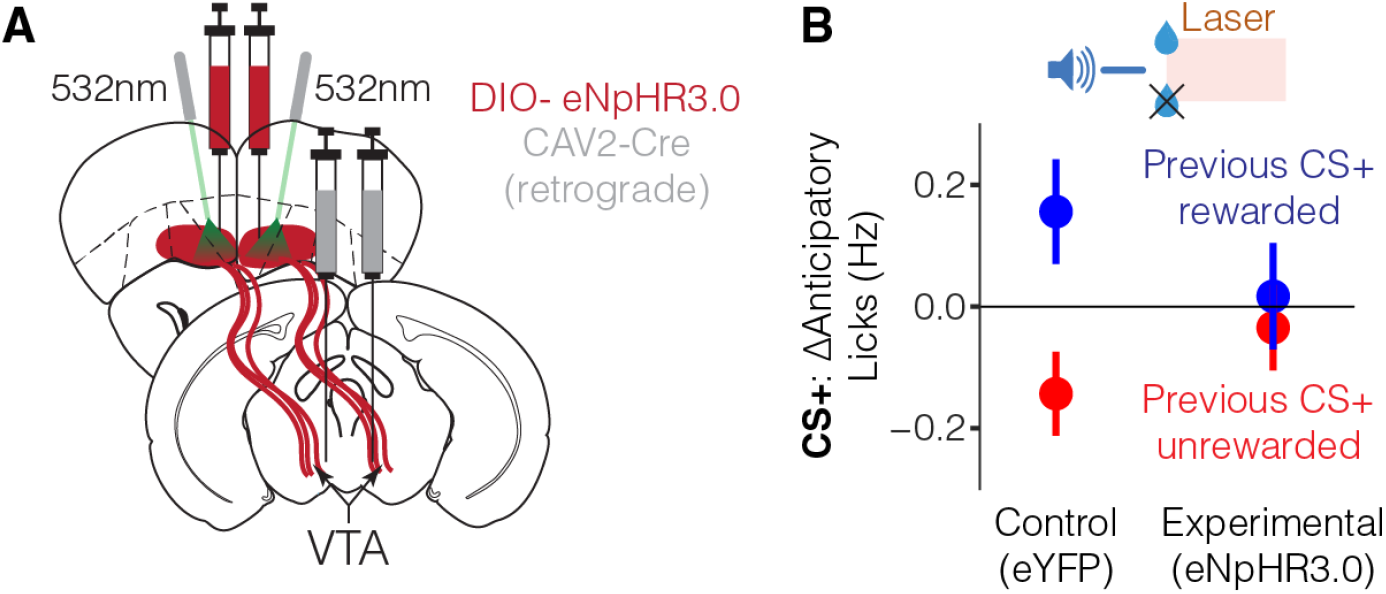
Trial-by-trial reward prediction learning is suppressed by the inhibition of vmOFC→VTA reward responses. **A.** Schematic for inhibition of vmOFC→VTA neurons. **B.** Results from experimental (those receiving inhibition) and control (those receiving no inhibition) animals showing abolishment of the trial-by-trial reward prediction learning due to inhibition. These data were previously published (Namboodiri et al., 2019) and are replotted here in the same format as Fig 1G. Statistical results are in the previous publication (Namboodiri et al., 2019).

**Fig S2 (related to Fig 2):**
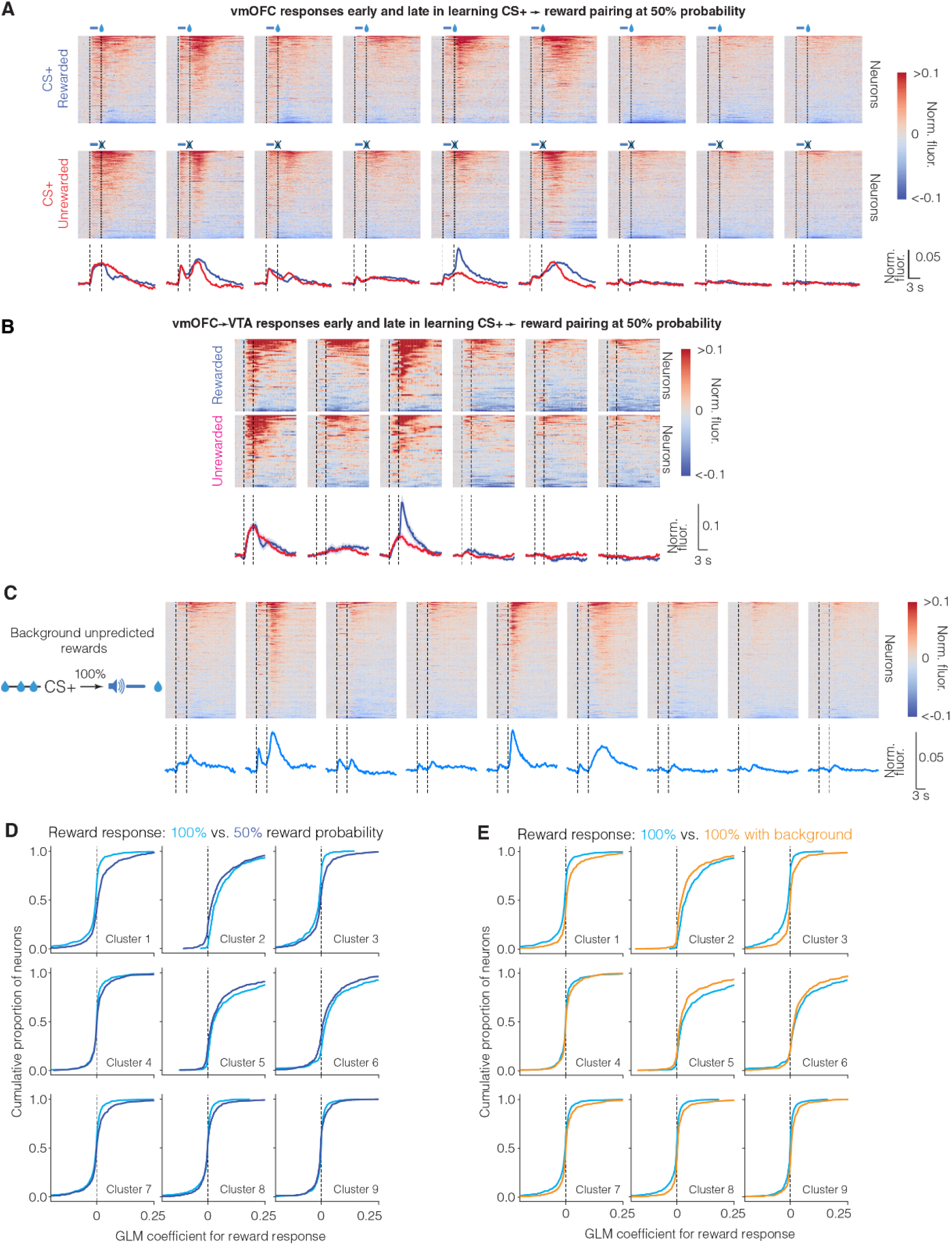
Characterization of vmOFC reward responses during two types of contingency degradation. **A.** PSTHs for all neurons in all clusters during the 50% probability session on rewarded and unrewarded trials. These data contain n=4,763 neurons from 5 mice. **B.** PSTHs of vmOFC→VTA neurons when reward probability was reduced to 50%. vmOFC→VTA neurons largely comprised of six of the clusters/subpopulations observed in vmOFC-CaMKII neurons. **C.** PSTHs of vmOFC CaMKIIα expressing neurons when reward probability was 100%, but random unpredictable rewards were delivered during the intertrial interval (“Background” session) (Namboodiri et al., 2019). The reward responses are positive in all clusters, much like they were early in learning. **D.** Cumulative distribution functions depicting the change in reward responses for all longitudinally tracked neurons after contingency degradation due to 50% probability reduction. The reward response on rewarded CS+ trials is compared between the two sessions. **E.** Cumulative distribution functions depicting the change in reward responses for all longitudinally tracked neurons after contingency degradation due to the presence of unpredictable rewards. Comparing **D** and **E** shows that both types of contingency degradations produce similar changes in reward responses. These data also directly argue against a model in which satiety or satiation affect the reward responses, as both a session that reduces the total number of rewards (50% reward probability) and a session that considerably increases the total number of rewards (background unpredictable rewards) produce similar changes to vmOFC reward responses (see **Supplementary Note 2**).

**Fig S3 (related to Fig 3):**
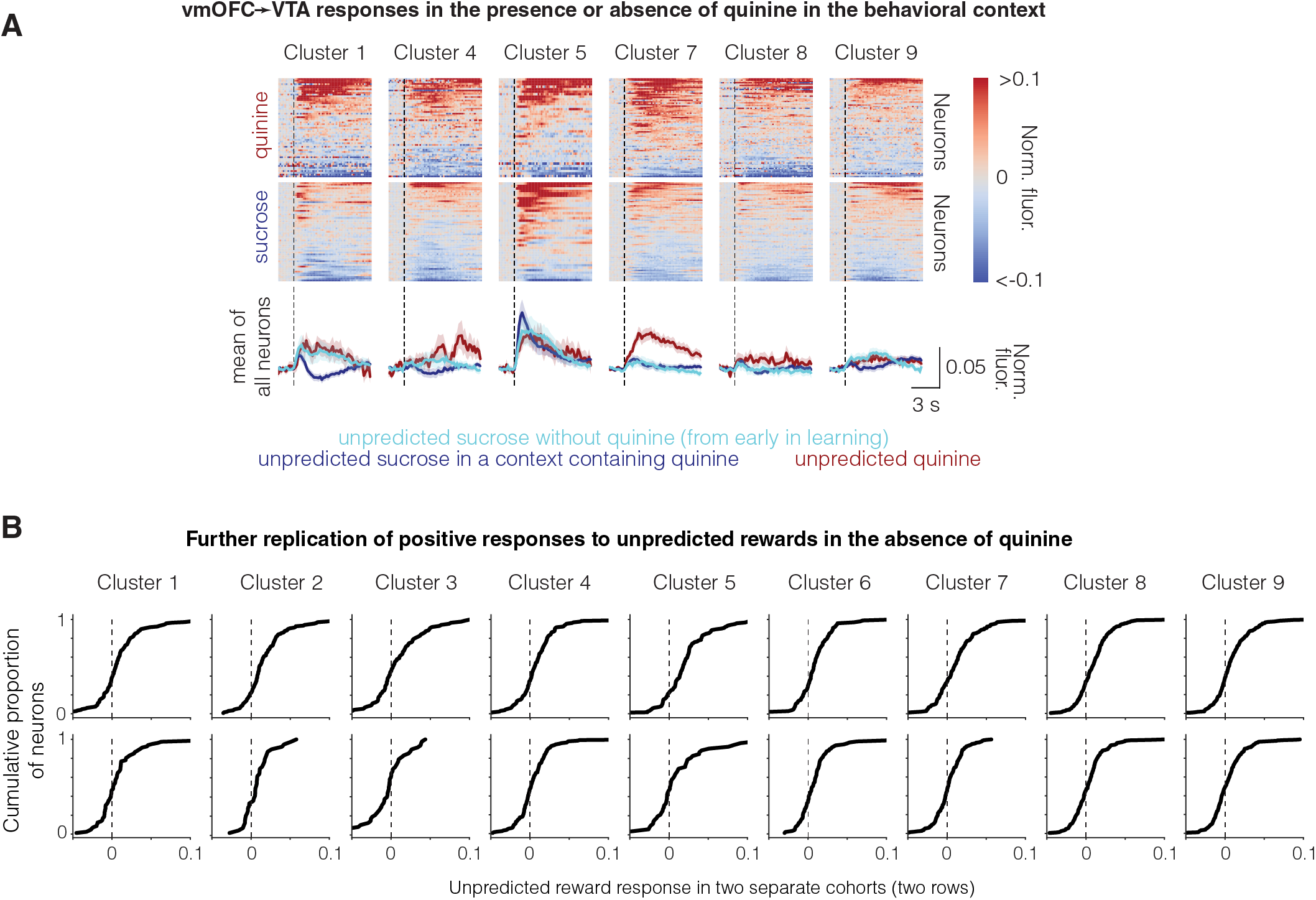
**A.** PSTH from all vmOFC→VTA neurons during sucrose and quinine, with responses from early in learning overlaid. Similar plot as in Fig 3D for vmOFC→VTA neurons. **B.** Unpredicted reward responses in a context without quinine in two separate cohorts of animals (same animals as those shown in Fig 6 and **Fig S6**, showing positive responses. Here, responses for 0-3 s after reward delivery in the absence of any LED presentation were quantified relative to pre-reward baseline.

**Fig S4 (related to Fig 4):**
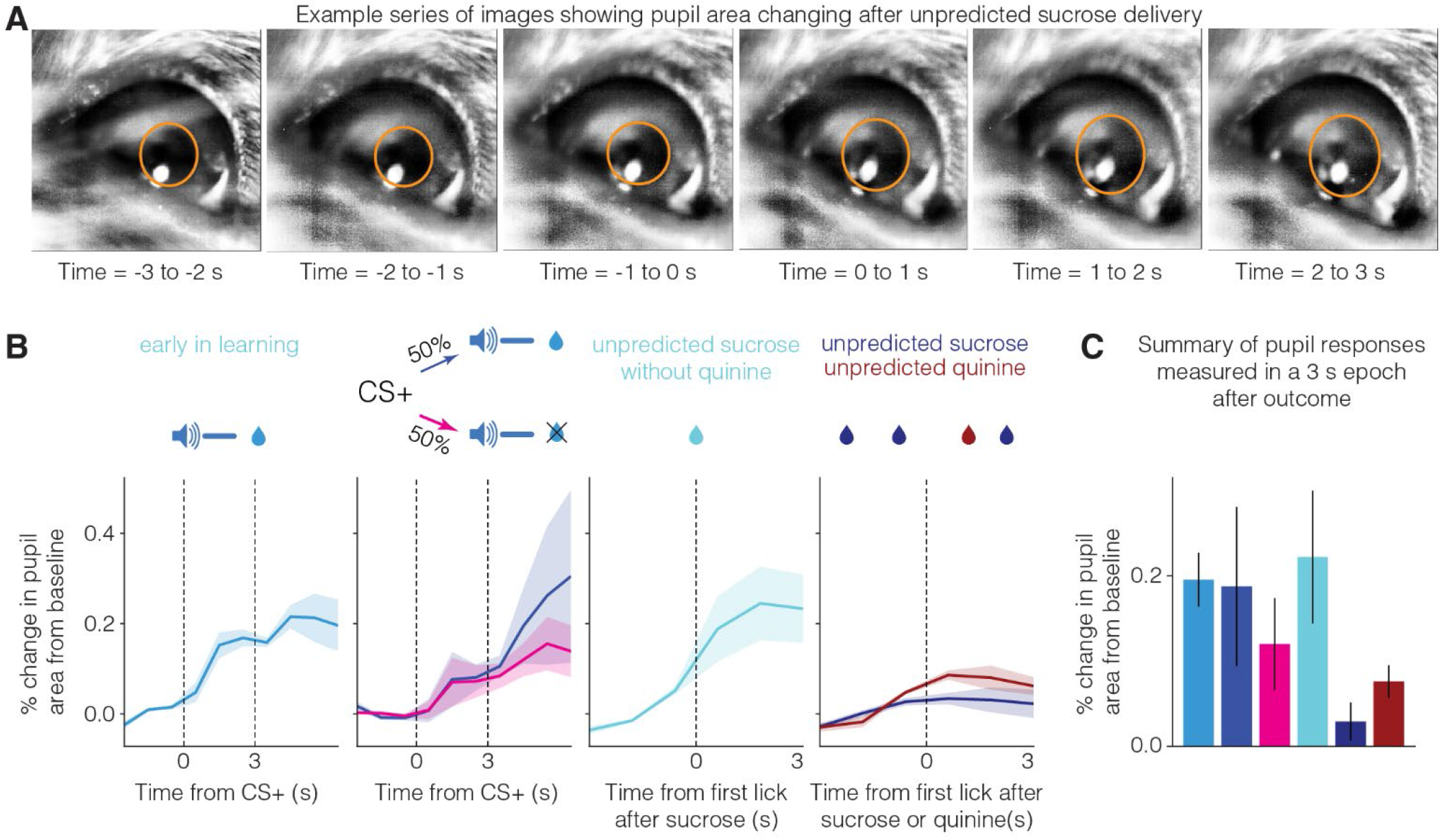
Changes in pupil diameter to reward across various task conditions. Example series of images of pupil dynamics one second apart each, showing an increase in pupil area during sucrose consumption. **B.** Dynamics of pupil responses in four task conditions. Pupil diameter was measured at 1 Hz resolution with smoothing (Methods). The elevated baseline immediately prior to first lick in the sucrose/quinine session likely reflects sensation of the delivered drop prior to the first lick. **C.** Summary of pupil area responses across the different conditions. Data were collected from n=4 animals each across all conditions. The responses early in learning and after 50% reward probability reduction do not correlate with the vmOFC reward responses shown in Fig 2 and **Fig S2**. Unpredicted sucrose in a context containing quinine produces a highly dampened pupil dilation response compared to unpredicted sucrose in a session without quinine.

**Fig S5 (related to Fig 5):**
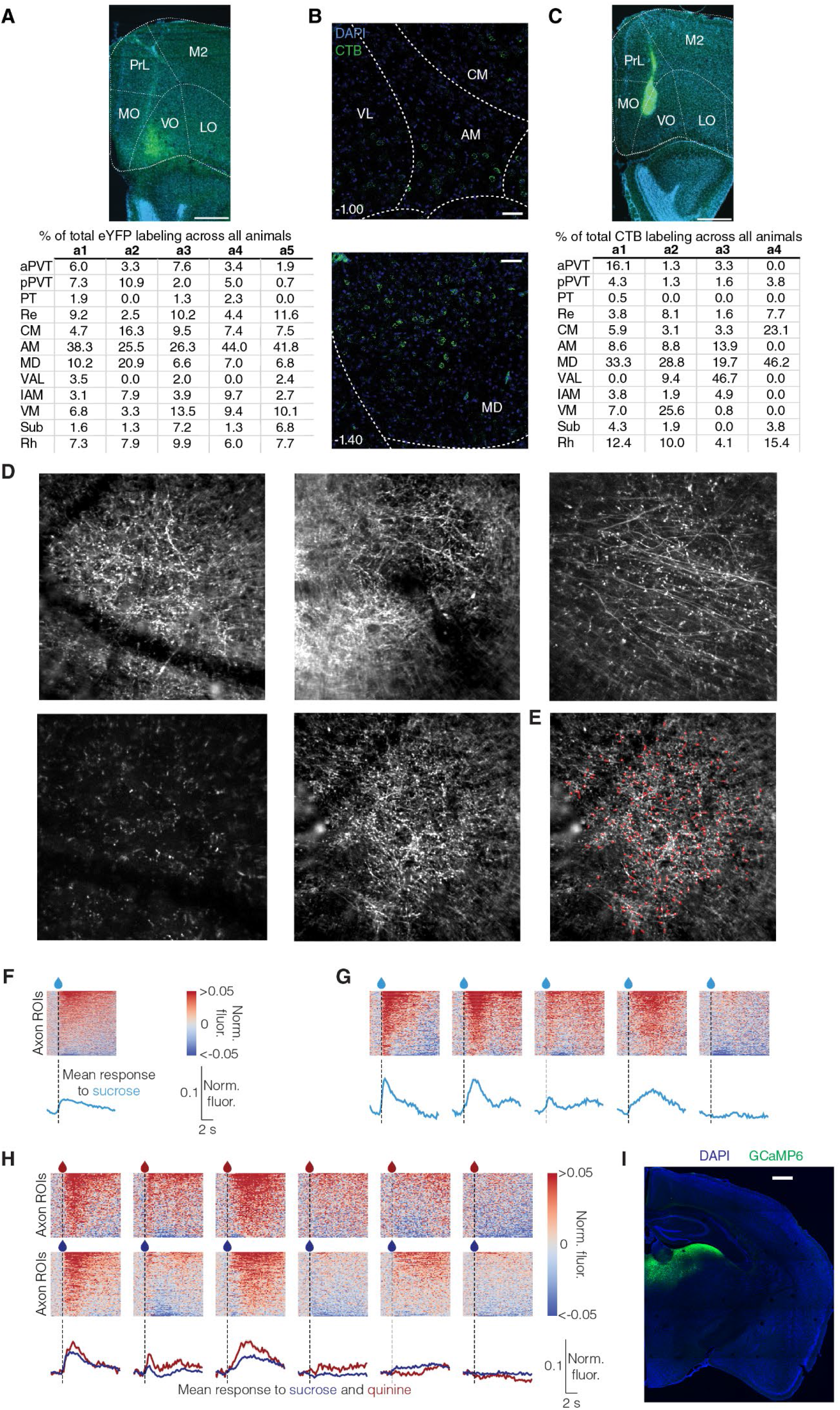
rAAV2retro and CTB injection sites, signal quantification, and heterogeneity in mThal→vmOFC axonal reward responses. **A.** (Top) Example injection site for rAAV2retro injection into the OFC for anatomical tracing of inputs to vmOFC. Scale bar = 500 µm. Blue = DAPI, green = eYFP. (Bottom) Table illustrating the breakdown of the percent of eYFP+ cells in each thalamic subregion of each animal (animals a1-a5). AM: Anteromedial, aPVT: Anterior paraventricular thalamus, Cg2: cingulate cortex area 2, CM: Centromedial, IAM: Interanteromedial, LO: lateral orbitofrontal cortex, M2: secondary motor cortex, MD: Mediodorsal, MO: medial orbitofrontal cortex, pPVT: Posterior Paraventricular thalamus, PrL: prelimbic cortex, PT: parataenial nucleus, Re: Nucleus of reuniens, Rh: Rhomboid nucleus, Sub: Submedial nucleus, VAL: Ventral anterior-lateral, VM: Ventromedial, VO: ventral orbitofrontal cortex. **B.** Representative images for CTB-488 labeling in the thalamus and quantification of labeling in animals receiving CTB-488 injection in the vmOFC. Scale bar 50 µm. **C.** Same as in **A** for CTB-488 injections. Since the difference between CTB and rAAV2retro labeling could be either due to viral tropism or due to slight differences in injection sites, we decided to pool these together to create an average map of mThal input regions to vmOFC in Fig 3B. **D.** Standard deviation projection images of activity from the entire fields of view (∼270 μm) of five different animals with mThal**→**vmOFC axons. Brightness and contrast were adjusted to maximize visibility. **E.** Manually drawn ROIs for individual axon segments shown in red. We took care to not double count different ROIs from the same axon (i.e. axons that extend within the imaging plane). However, since axons branch in and out of the imaging focal plane, it is possible that different axonal ROIs actually derive from the same axon. We did not attempt to classify ROIs as resulting from a single axon using a threshold of activity correlation since such measures depend crucially on the signal to noise ratio of the recording; for low signal to noise ratio recordings, two segments from the exact same axon can have low correlation. Thus, we instead only make claims about the population activity, either as the average, or as the average of an activity-defined cluster (**G**, **H**). **F.** Same as in Fig 3F but including all five animals. The sucrose quinine experiment could not be run in the first two animals shown in the top row. **G.** Results of clustering axonal activity patterns (same approach as used for vmOFC cell bodies) showing five clusters of axon ROIs with distinct response profiles to sucrose. This analysis shows that distinct reward response profiles exist within mThal**→**vmOFC axons. The benefit of analyzing using a clustering approach is that we know that axon ROIs that are potentially from the same axons have to necessarily be within the same cluster since they will show the same response pattern (i.e. signal is the same irrespective of signal to noise ratio). Thus, we only interpret the results at the level of a cluster and state that these distinct response profiles exist within mThal**→**vmOFC axons. An important caveat is that we cannot make any claims regarding the relative prevalence of each response type or cluster due to the potential of double counting of axons within ROIs. **H.** Same as in **G** but for sucrose and quinine responses. The clustering was done separately for all axon ROIs recorded in this session as they were not longitudinally tracked from the sucrose only session. The results qualitatively match response patterns in vmOFC since 4 out of 6 clusters show larger responses for quinine compared to sucrose. Interestingly, two clusters (right most) appear to have larger responses for sucrose compared to quinine, potentially signaling value or valence of the liquid. **I.** Representative histological image showing expression of GCaMP6s in mThal cell bodies. Though there is relatively widespread expression in thalamus, only axons projecting to vmOFC were imaged (shown in **D**). Scale bar is 500 μm.

**Fig S6 (related to Fig 6):**
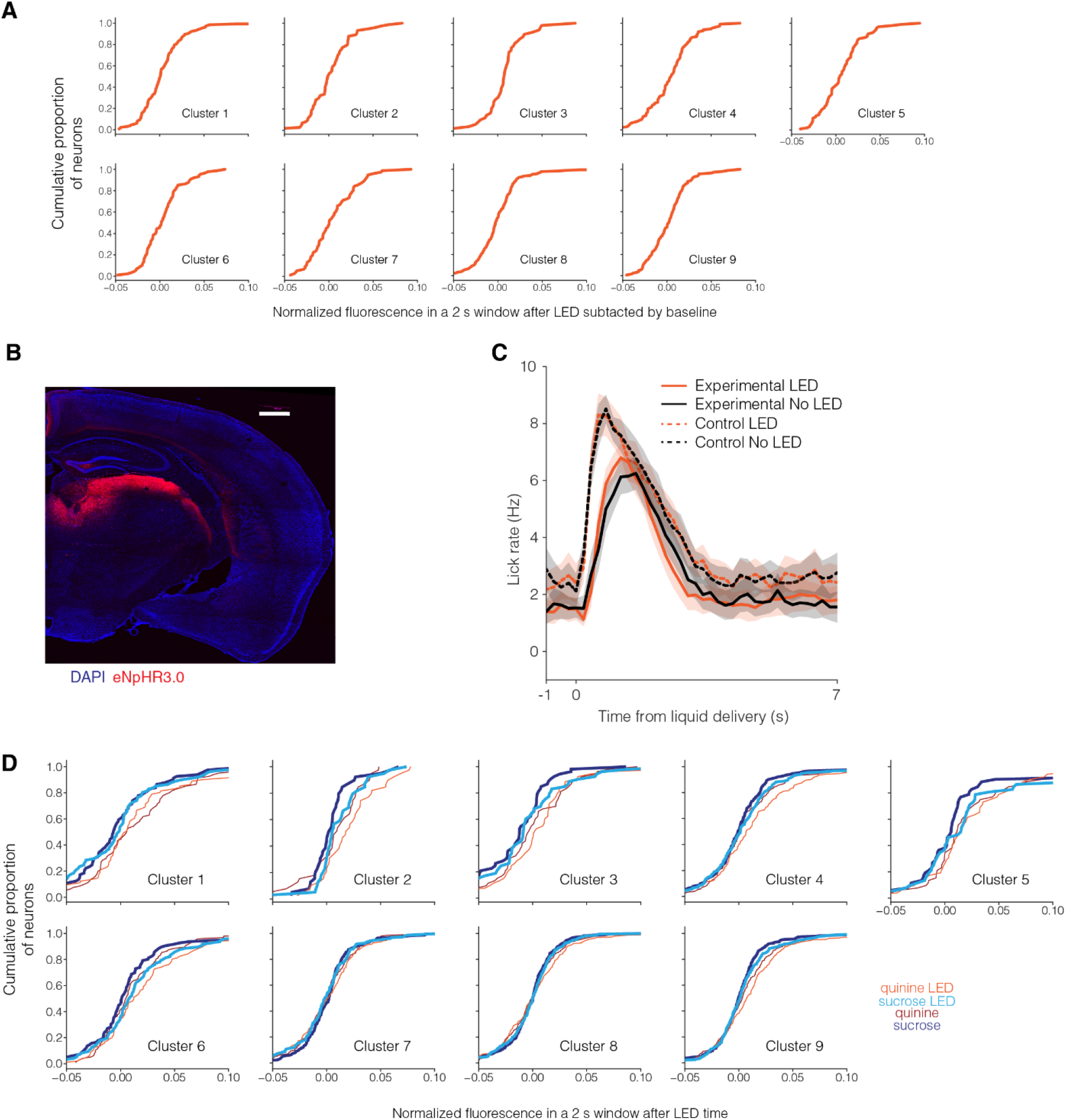
Controls for mThal→vmOFC axon inhibition. **A.** Empirical cumulative distribution function of vmOFC neuronal responses with LED in experimental animals expressing eNpHR3.0, but in the absence of any rewards (i.e. spontaneous inhibition of inputs). The responses were compared to the baseline right before inhibition. No cluster had a mean LED response significantly different from zero. **B.** Representative histological image showing expression of eNpHR3.0-mCherry in mThal cell bodies. Though there is relatively widespread expression in thalamus, only axons projecting to vmOFC were inhibited. Scale bar is 500 μm. **C.** Average licking behavior across trials with or without LED in experimental and control mThal inhibition groups. There was no difference in licking due to the presence of LED in either group. **D.** Empirical cumulative distribution function of vmOFC neuronal responses to sucrose and quinine with and without LED in control animals expressing a fluorophore (mCherry) without eNpHR3.0. Clusters 5 and 6 had a mean LED response significantly different from zero for sucrose, and clusters 2, 4, 6 and 9 had a mean LED response significantly different from zero for quinine. This suggests that there is a non-selective change in quinine responses in some clusters (as also seen in clusters 1 and 4 in Fig 6). The sucrose response changed due to LED in 7 out of 9 clusters in experimental animals and 2 out of 9 in the controls, likely suggesting that most of it is a selective change due to inhibition of the inputs. A direct comparison between the two groups is not possible since the expression levels of GCaMP6s and eNpHR3.0 vary across animals.

**Table S1.**
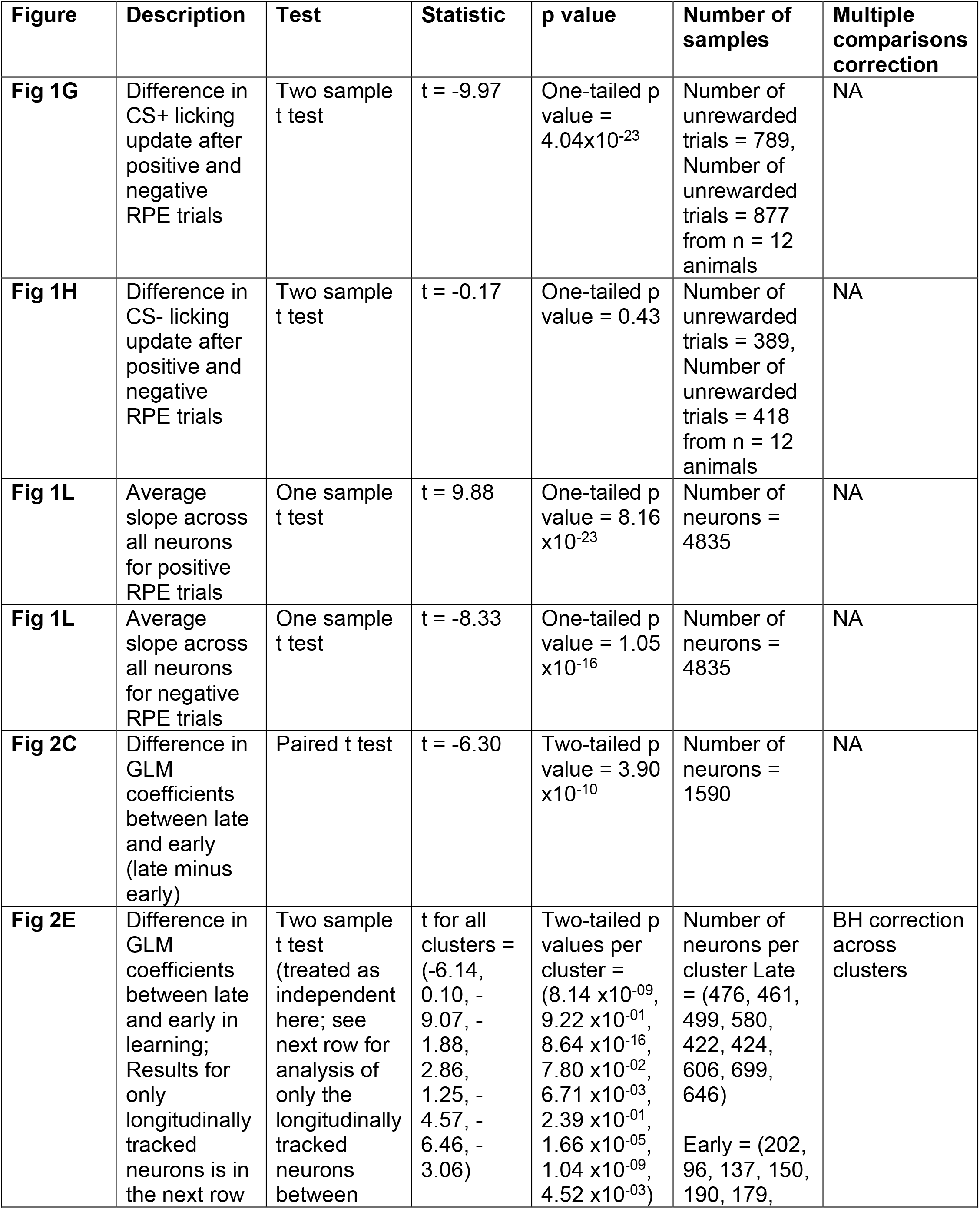
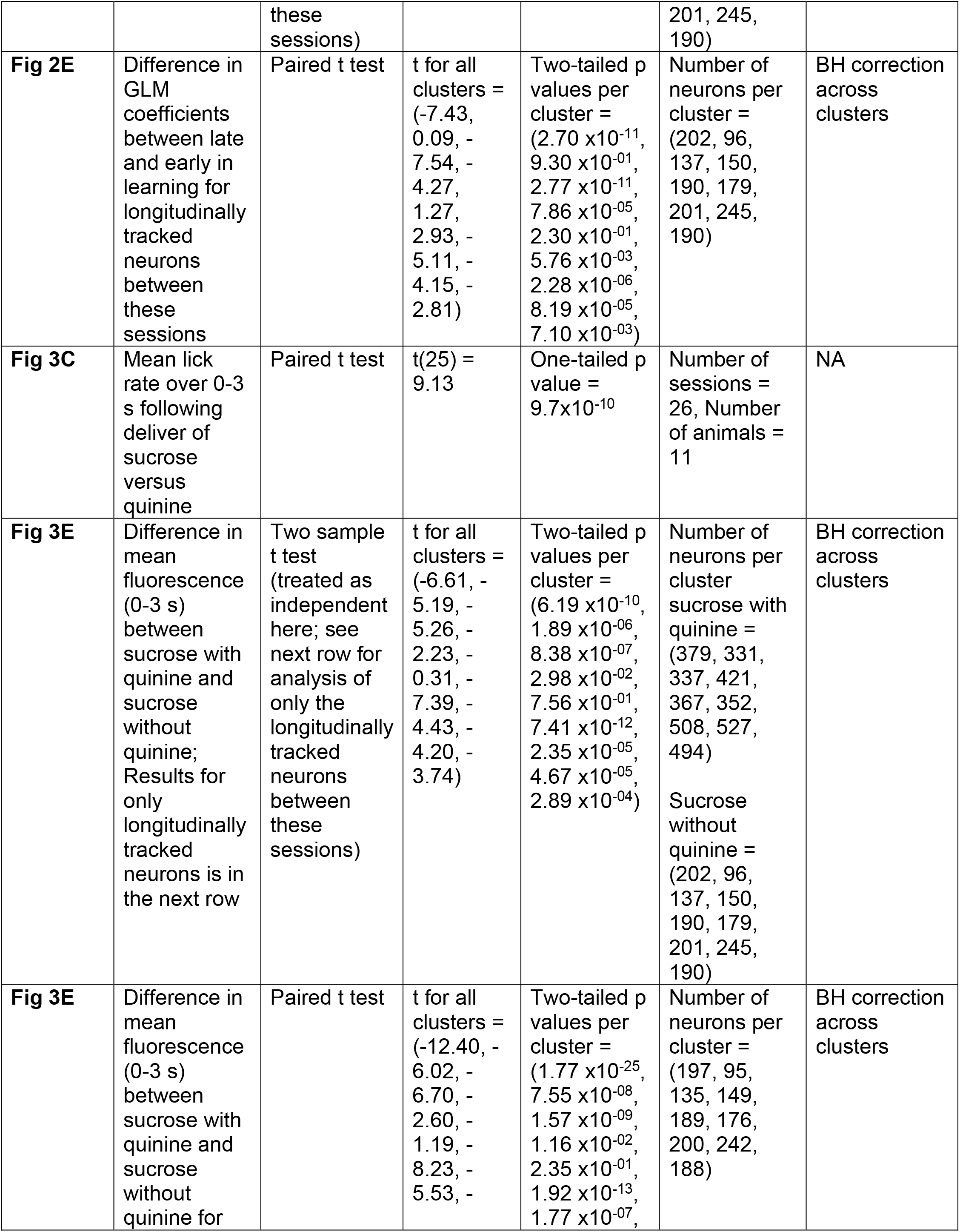
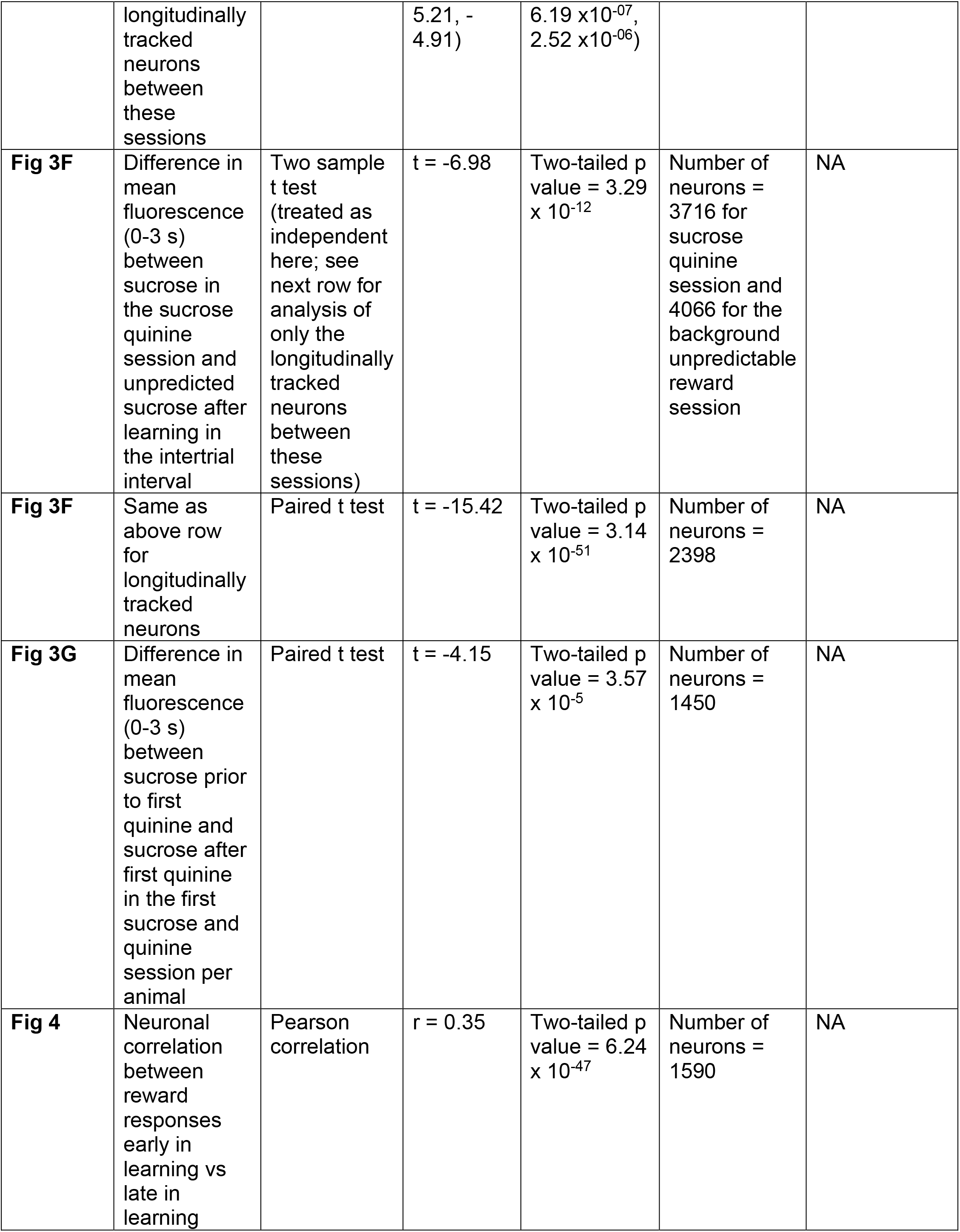
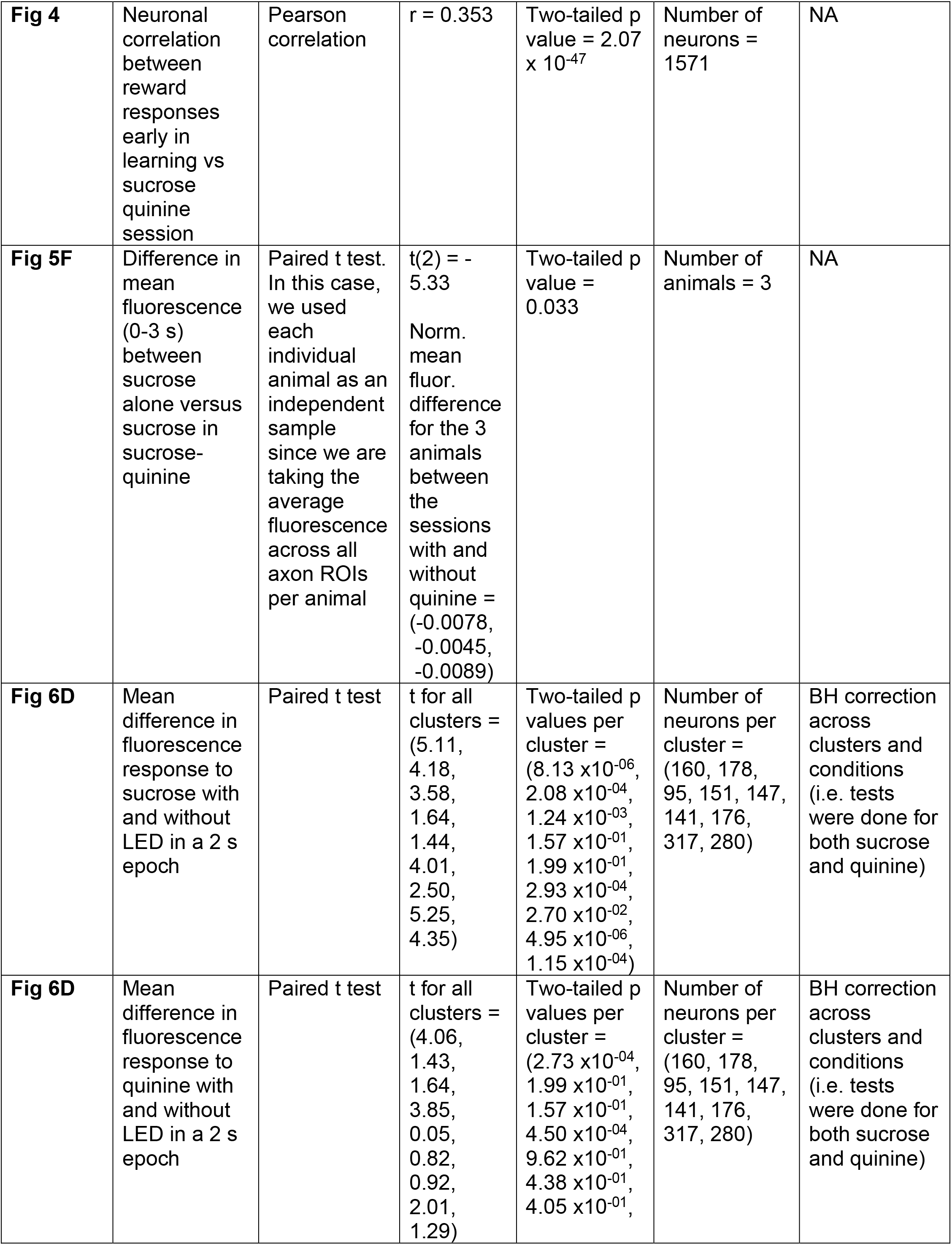
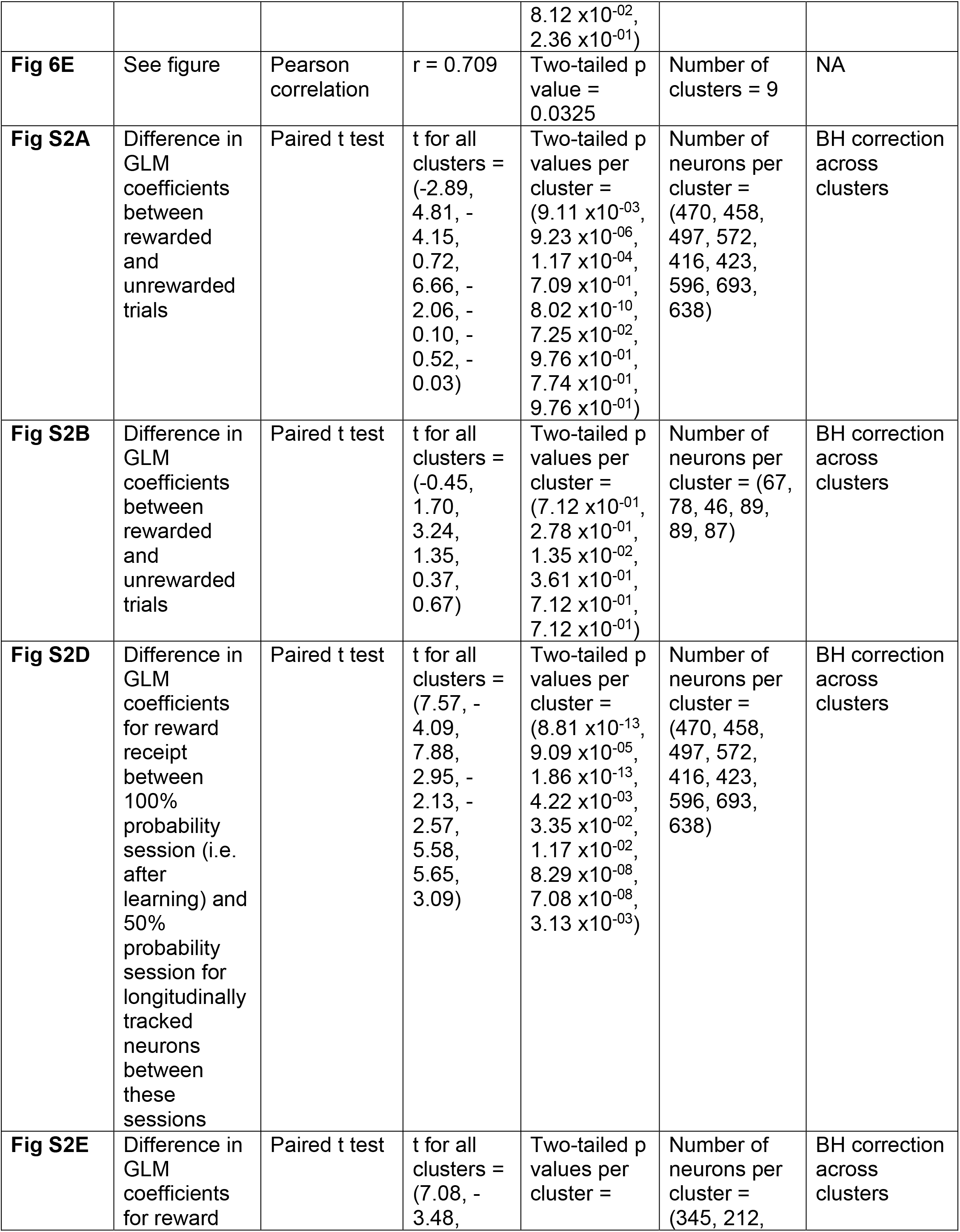
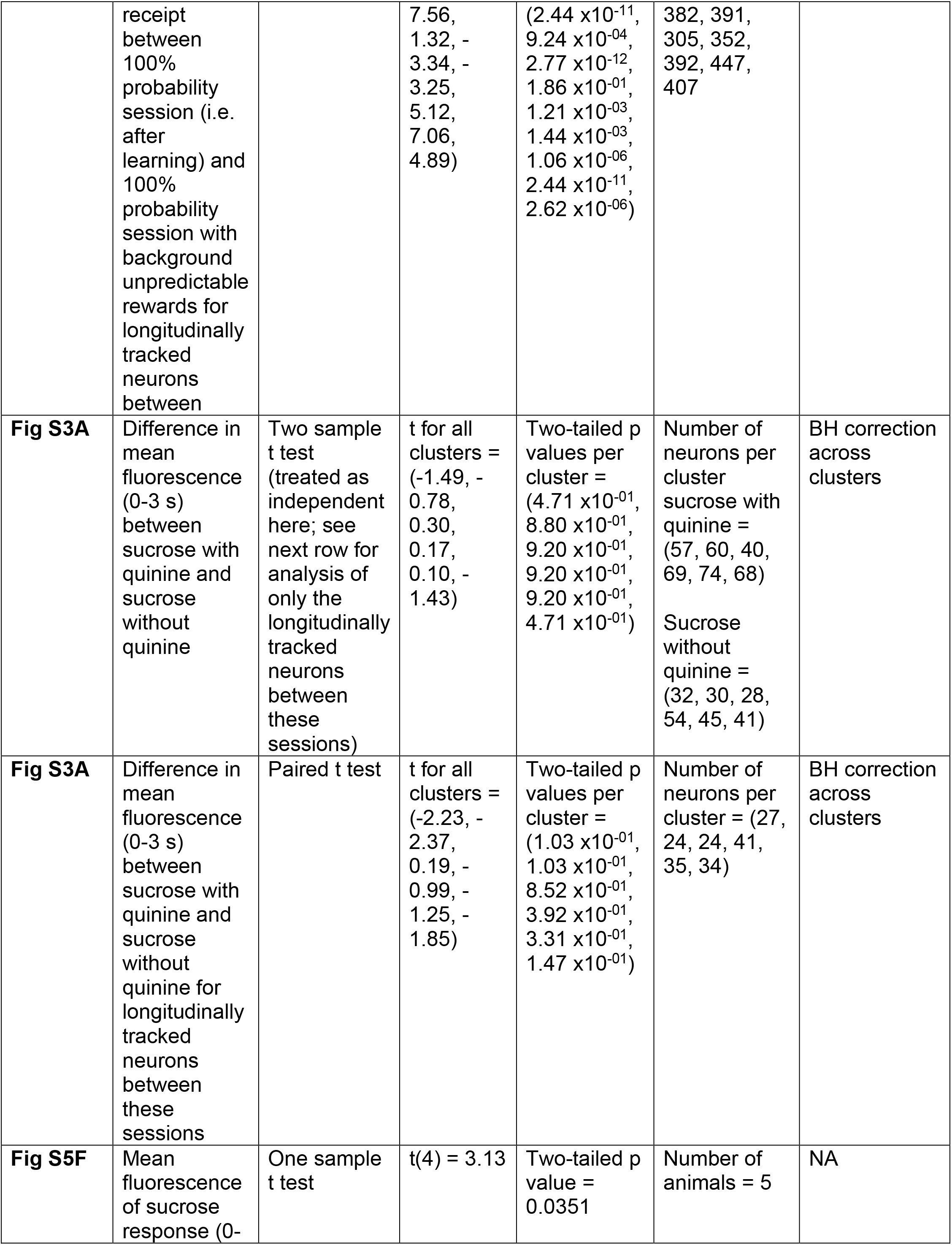
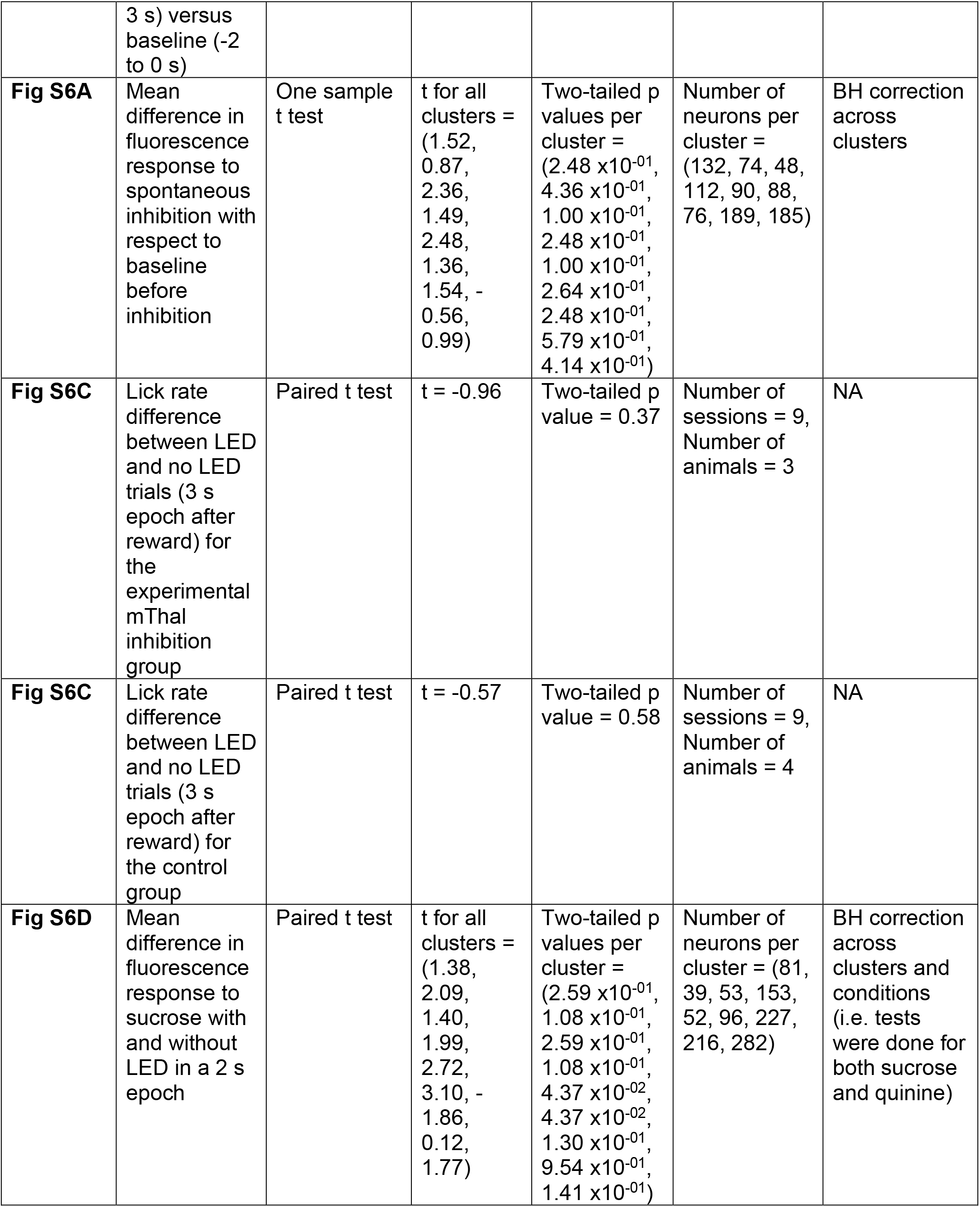
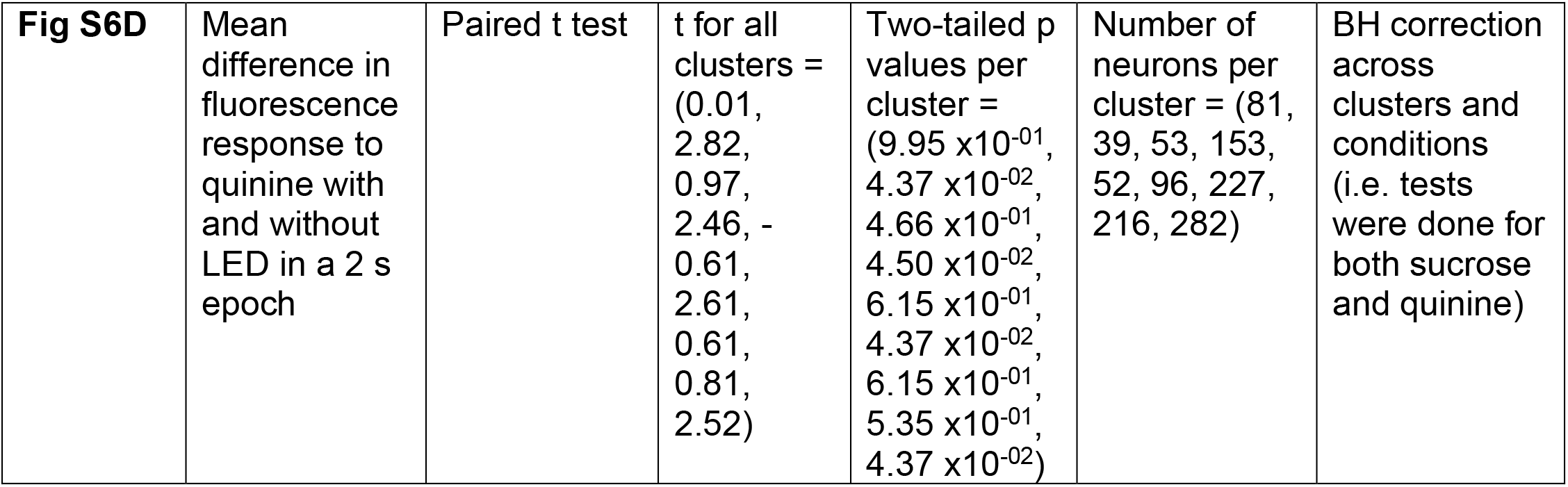
(Statistical Results)

## References

Ader, R., Weijnen, J.A.W.M., and Moleman, P. (1972). Retention of a passive avoidance response as a function of the intensity and duration of electric shock. Psychon Sci 26, 125–128.

Amarante, L.M., and Laubach, M. (2020). Rhythmic activity in the medial and orbital frontal cortices tracks reward value and the vigor of consummatory behavior. BioRxiv 2020.09.22.308809.

Banerjee, A., Parente, G., Teutsch, J., Lewis, C., Voigt, F.F., and Helmchen, F. (2020). Value-guided remapping of sensory cortex by lateral orbitofrontal cortex. Nature 585, 245–250.

Behrens, T.E.J., Woolrich, M.W., Walton, M.E., and Rushworth, M.F.S. (2007). Learning the value of information in an uncertain world. Nat Neurosci 10, 1214–1221.

Bower, G.H., and Trabasso, T. (1964). Concept Identification. In Studies in Mathematical Psychology, R.C. Atkinson, ed. (Stanford University Press), p.

Bradfield, L.A., and Hart, G. (2020). Rodent medial and lateral orbitofrontal cortices represent unique components of cognitive maps of task space. Neurosci Biobehav Rev 108, 287–294.

Chang, C.Y., Esber, G.R., Marrero-Garcia, Y., Yau, H.-J., Bonci, A., and Schoenbaum, G. (2016). Brief optogenetic inhibition of dopamine neurons mimics endogenous negative reward prediction errors. Nat Neurosci 19, 111–116.

Chen, T.-W., Wardill, T.J., Sun, Y., Pulver, S.R., Renninger, S.L., Baohan, A., Schreiter, E.R., Kerr, R.A., Orger, M.B., Jayaraman, V., et al. (2013). Ultrasensitive fluorescent proteins for imaging neuronal activity. Nature 499, 295–300.

Coddington, L.T., and Dudman, J.T. (2018). The timing of action determines reward prediction signals in identified midbrain dopamine neurons. Nat. Neurosci. 21, 1563–1573.

Cohen, J.Y., Haesler, S., Vong, L., Lowell, B.B., and Uchida, N. (2012). Neuron-type-specific signals for reward and punishment in the ventral tegmental area. Nature 482, 85–88.

Constantinople, C.M., Piet, A.T., Bibawi, P., Akrami, A., Kopec, C., and Brody, C.D. (2019). Lateral orbitofrontal cortex promotes trial-by-trial learning of risky, but not spatial, biases. Elife 8.

Costa, V.D., and Averbeck, B.B. (2020). Primate Orbitofrontal Cortex Codes Information Relevant for Managing Explore–Exploit Tradeoffs. J. Neurosci. 40, 2553–2561.

Courville, A.C., Daw, N.D., and Touretzky, D.S. (2006). Bayesian theories of conditioning in a changing world. Trends in Cognitive Sciences 10, 294–300.

Dabney, W., Kurth-Nelson, Z., Uchida, N., Starkweather, C.K., Hassabis, D., Munos, R., and Botvinick, M. (2020). A distributional code for value in dopamine-based reinforcement learning. Nature 577, 671–675.

Downing, B.D. (1968). Salience and learning rate in concept identification. Psychon Sci 10, 73–74.

Engelhard, B., Finkelstein, J., Cox, J., Fleming, W., Jang, H.J., Ornelas, S., Koay, S.A., Thiberge, S.Y., Daw, N.D., Tank, D.W., et al. (2019). Specialized coding of sensory, motor and cognitive variables in VTA dopamine neurons. Nature 570, 509–513.

Eshel, N., Bukwich, M., Rao, V., Hemmelder, V., Tian, J., and Uchida, N. (2015). Arithmetic and local circuitry underlying dopamine prediction errors. Nature 525, 243–246.

Everitt, B.J., Hutcheson, D.M., Ersche, K.D., Pelloux, Y., Dalley, J.W., and Robbins, T.W. (2007). The orbital prefrontal cortex and drug addiction in laboratory animals and humans. Ann. N. Y. Acad. Sci. 1121, 576–597.

Frank, M.J., Doll, B.B., Oas-Terpstra, J., and Moreno, F. (2009). Prefrontal and striatal dopaminergic genes predict individual differences in exploration and exploitation. Nat Neurosci 12, 1062–1068.

Galea, J.M., Mallia, E., Rothwell, J., and Diedrichsen, J. (2015). The dissociable effects of punishment and reward on motor learning. Nat. Neurosci. 18, 597–602.

Gavornik, J.P., Shuler, M.G.H., Loewenstein, Y., Bear, M.F., and Shouval, H.Z. (2009). Learning reward timing in cortex through reward dependent expression of synaptic plasticity. PNAS 106, 6826– 6831.

Gershman, S.J. (2015). Do learning rates adapt to the distribution of rewards? Psychon Bull Rev 22, 1320–1327.

Gremel, C.M., Chancey, J.H., Atwood, B.K., Luo, G., Neve, R., Ramakrishnan, C., Deisseroth, K., Lovinger, D.M., and Costa, R.M. (2016). Endocannabinoid Modulation of Orbitostriatal Circuits Gates Habit Formation. Neuron 90, 1312–1324.

Groman, S.M., Keistler, C., Keip, A.J., Hammarlund, E., DiLeone, R.J., Pittenger, C., Lee, D., and Taylor, J.R. (2019). Orbitofrontal Circuits Control Multiple Reinforcement-Learning Processes. Neuron 0.

Grossman, C.D., Bari, B.A., and Cohen, J.Y. (2020). Serotonin neurons modulate learning rate through uncertainty. BioRxiv 2020.10.24.353508.

Halassa, M.M., and Sherman, S.M. (2019). Thalamocortical Circuit Motifs: A General Framework. Neuron 103, 762–770.

Hayden, B.Y., Heilbronner, S.R., Pearson, J.M., and Platt, M.L. (2011). Surprise Signals in Anterior Cingulate Cortex: Neuronal Encoding of Unsigned Reward Prediction Errors Driving Adjustment in Behavior. J Neurosci 31, 4178–4187.

Hirokawa, J., Vaughan, A., Masset, P., Ott, T., and Kepecs, A. (2019). Frontal cortex neuron types categorically encode single decision variables. Nature 576, 446–451.

Holly, E.N., Davatolhagh, M.F., Choi, K., Alabi, O.O., Vargas Cifuentes, L., and Fuccillo, M.V. (2019). Striatal Low-Threshold Spiking Interneurons Regulate Goal-Directed Learning. Neuron 103, 92–101.e6.

Iigaya, K. (2016). Adaptive learning and decision-making under uncertainty by metaplastic synapses guided by a surprise detection system. ELife 5, e18073.

Izquierdo, A., and Murray, E.A. (2010). Functional interaction of medial MD thalamus but not nucleus accumbens with amygdala and orbital prefrontal cortex is essential for adaptive response selection after reinforcer devaluation. J Neurosci 30, 661–669.

Jankowski, M.M., Ronnqvist, K.C., Tsanov, M., Vann, S.D., Wright, N.F., Erichsen, J.T., Aggleton, J.P., and O’Mara, S.M. (2013). The anterior thalamus provides a subcortical circuit supporting memory and spatial navigation. Front Syst Neurosci 7.

Jennings, J.H., Kim, C.K., Marshel, J.H., Raffiee, M., Ye, L., Quirin, S., Pak, S., Ramakrishnan, C., and Deisseroth, K. (2019). Interacting neural ensembles in orbitofrontal cortex for social and feeding behaviour. Nature 565, 645–649.

Jones, J.L., Esber, G.R., McDannald, M.A., Gruber, A.J., Hernandez, A., Mirenzi, A., and Schoenbaum, G. (2012). Orbitofrontal Cortex Supports Behavior and Learning Using Inferred But Not Cached Values. Science 338, 953–956.

Kaifosh, P., Zaremba, J.D., Danielson, N.B., and Losonczy, A. (2014). SIMA: Python software for analysis of dynamic fluorescence imaging data. Front Neuroinform 8, 80.

Kepecs, A., Uchida, N., Zariwala, H.A., and Mainen, Z.F. (2008). Neural correlates, computation and behavioural impact of decision confidence. Nature 455, 227–231.

Kheifets, A., Freestone, D., and Gallistel, C.R. (2017). THEORETICAL IMPLICATIONS OF QUANTITATIVE PROPERTIES OF INTERVAL TIMING AND PROBABILITY ESTIMATION IN MOUSE AND RAT. J Exp Anal Behav 108, 39–72.

Kobayashi, S., and Schultz, W. (2008). Influence of reward delays on responses of dopamine neurons. J. Neurosci. 28, 7837–7846.

Kojima, S., Yamanaka, M., Fujito, Y., and Ito, E. (1996). Differential Neuroethological Effects of Aversive and Appetitive Reinforcing Stimuli on Associative Learning in Lymnaea stagnalis. Jzoo 13, 803–812.

Lee, K., Claar, L.D., Hachisuka, A., Bakhurin, K.I., Nguyen, J., Trott, J.M., Gill, J.L., and Masmanidis, S.C. (2020). Temporally restricted dopaminergic control of reward-conditioned movements. Nat. Neurosci. 23, 209–216.

Louie, K., Grattan, L.E., and Glimcher, P.W. (2011). Reward Value-Based Gain Control: Divisive Normalization in Parietal Cortex. J. Neurosci. 31, 10627–10639.

Mackintosh, N.J. (1976). Overshadowing and stimulus intensity. Animal Learning & Behavior 4, 186–192.

Matsumoto, H., Tian, J., Uchida, N., and Watabe-Uchida, M. (2016). Midbrain dopamine neurons signal aversion in a reward-context-dependent manner. Elife 5.

Miller, K.J., Botvinick, M.M., and Brody, C.D. (2018). Value Representations in Orbitofrontal Cortex Drive Learning, but not Choice. BioRxiv 245720.

Mitchell, A.S., and Chakraborty, S. (2013). What does the mediodorsal thalamus do? Front Syst Neurosci 7, 37.

Mohebi, A., Pettibone, J.R., Hamid, A.A., Wong, J.-M.T., Vinson, L.T., Patriarchi, T., Tian, L., Kennedy, R.T., and Berke, J.D. (2019). Dissociable dopamine dynamics for learning and motivation. Nature 570, 65–70.

Monosov, I.E., and Hikosaka, O. (2012). Regionally Distinct Processing of Rewards and Punishments by the Primate Ventromedial Prefrontal Cortex. J. Neurosci. 32, 10318–10330.

Morisot, N., Phamluong, K., Ehinger, Y., Berger, A.L., Moffat, J.J., and Ron, D. (2019). mTORC1 in the orbitofrontal cortex promotes habitual alcohol seeking. Elife 8.

Namboodiri, V.M.K., Huertas, M.A., Monk, K.J., Shouval, H.Z., and Hussain Shuler, M.G. (2015). Visually cued action timing in the primary visual cortex. Neuron 86, 319–330.

Namboodiri, V.M.K., Otis, J.M., Heeswijk, K. van, Voets, E.S., Alghorazi, R.A., Rodriguez-Romaguera, J., Mihalas, S., and Stuber, G.D. (2019). Single-cell activity tracking reveals that orbitofrontal neurons acquire and maintain a long-term memory to guide behavioral adaptation. Nat. Neurosci. 22, 1110.

Otis, J.M., Namboodiri, V.M.K., Matan, A.M., Voets, E.S., Mohorn, E.P., Kosyk, O., McHenry, J.A., Robinson, J.E., Resendez, S.L., Rossi, M.A., et al. (2017). Prefrontal cortex output circuits guide reward seeking through divergent cue encoding. Nature 543, 103–107.

Otis, J.M., Zhu, M., Namboodiri, V.M.K., Cook, C.A., Kosyk, O., Matan, A.M., Ying, R., Hashikawa, Y., Hashikawa, K., Trujillo-Pisanty, I., et al. (2019). Paraventricular Thalamus Projection Neurons Integrate Cortical and Hypothalamic Signals for Cue-Reward Processing. Neuron 103, 423–431.e4.

Ottenheimer, D., Richard, J.M., and Janak, P.H. (2018). Ventral pallidum encodes relative reward value earlier and more robustly than nucleus accumbens. Nature Communications 9, 4350.

Padoa-Schioppa, C., and Assad, J.A. (2006). Neurons in the orbitofrontal cortex encode economic value. Nature 441, 223–226.

Padoa-Schioppa, C., and Assad, J.A. (2008). The representation of economic value in the orbitofrontal cortex is invariant for changes of menu. Nat. Neurosci. 11, 95–102.

Parent, M.A., Amarante, L.M., Liu, B., Weikum, D., and Laubach, M. (2015). The medial prefrontal cortex is crucial for the maintenance of persistent licking and the expression of incentive contrast. Front Integr Neurosci 9, 23.

Pascoli, V., Hiver, A., Van Zessen, R., Loureiro, M., Achargui, R., Harada, M., Flakowski, J., and Lüscher, C. (2018). Stochastic synaptic plasticity underlying compulsion in a model of addiction. Nature 564, 366–371.

Pearce, J.M., and Hall, G. (1980). A model for Pavlovian learning: variations in the effectiveness of conditioned but not of unconditioned stimuli. Psychol Rev 87, 532–552.

Preuschoff, K., and Bossaerts, P. (2007). Adding prediction risk to the theory of reward learning. Ann. N. Y. Acad. Sci. 1104, 135–146.

Rescorla, R.A., and Wagner, A.R. (1972). A theory of Pavlovian conditioning: Variations in the effectiveness of reinforcement and nonreinforcement. Classical Conditioning II: Current Research and Theory 2, 64–99.

Resendez, S.L., Jennings, J.H., Ung, R.L., Namboodiri, V.M.K., Zhou, Z.C., Otis, J.M., Nomura, H., McHenry, J.A., Kosyk, O., and Stuber, G.D. (2016). Visualization of cortical, subcortical and deep brain neural circuit dynamics during naturalistic mammalian behavior with head-mounted microscopes and chronically implanted lenses. Nat Protoc 11, 566–597.

Roesch, M.R., Esber, G.R., Li, J., Daw, N.D., and Schoenbaum, G. (2012). Surprise! Neural correlates of Pearce-Hall and Rescorla-Wagner coexist within the brain. Eur. J. Neurosci. 35, 1190– 1200.

Schultz, W., Dayan, P., and Montague, P.R. (1997). A Neural Substrate of Prediction and Reward. Science 275, 1593–1599.

Schweighofer, N., and Doya, K. (2003). Meta-learning in reinforcement learning. Neural Netw 16, 5–9.

Simmons, J.M., and Richmond, B.J. (2008). Dynamic changes in representations of preceding and upcoming reward in monkey orbitofrontal cortex. Cereb Cortex 18, 93–103.

Slotnick, B., and Coppola, D.M. (2015). Odor-Cued Taste Avoidance: A Simple and Robust Test of Mouse Olfaction. Chem Senses 40, 269–278.

Soltani, A., and Izquierdo, A. (2019). Adaptive learning under expected and unexpected uncertainty. Nat Rev Neurosci 20, 635–644.

Steinberg, E.E., Keiflin, R., Boivin, J.R., Witten, I.B., Deisseroth, K., and Janak, P.H. (2013). A causal link between prediction errors, dopamine neurons and learning. Nat Neurosci 16, 966–973.

Sutton, R.S., and Barto, A.G. (1998). Introduction to Reinforcement Learning (Cambridge, MA, USA: MIT Press).

Tervo, D.G.R., Hwang, B.-Y., Viswanathan, S., Gaj, T., Lavzin, M., Ritola, K.D., Lindo, S., Michael, S., Kuleshova, E., Ojala, D., et al. (2016). A Designer AAV Variant Permits Efficient Retrograde Access to Projection Neurons. Neuron 92, 372–382.

Tremblay, L., and Schultz, W. (1999). Relative reward preference in primate orbitofrontal cortex. Nature 398, 704–708.

Wang, J.X., Kurth-Nelson, Z., Kumaran, D., Tirumala, D., Soyer, H., Leibo, J.Z., Hassabis, D., and Botvinick, M. (2018). Prefrontal cortex as a meta-reinforcement learning system. Nat. Neurosci. 21, 860–868.

Wang, P.Y., Boboila, C., Chin, M., Higashi-Howard, A., Shamash, P., Wu, Z., Stein, N.P., Abbott, L.F., and Axel, R. (2020). Transient and Persistent Representations of Odor Value in Prefrontal Cortex. Neuron 108, 209–224.e6.

Wilson, R.C., Takahashi, Y.K., Schoenbaum, G., and Niv, Y. (2014). Orbitofrontal cortex as a cognitive map of task space. Neuron 81, 267–279.

Zhong, W., Li, Y., Feng, Q., and Luo, M. (2017). Learning and Stress Shape the Reward Response Patterns of Serotonin Neurons. J. Neurosci. 37, 8863–8875.

Zuiderveld, K. (1994). Contrast limited adaptive histogram equalization. In Graphics Gems IV, (USA: Academic Press Professional, Inc.), pp. 474–485.

